# Generation of diversity in the blue cheese mold *Penicillium roqueforti* and identification of pleiotropic QTL for key cheese-making phenotypes

**DOI:** 10.1101/2024.02.22.581506

**Authors:** Thibault Caron, Ewen Crequer, Mélanie Le Piver, Stéphanie Le Prieur, Sammy Brunel, Alodie Snirc, Gwennina Cueff, Daniel Roueyre, Michel Place, Christophe Chassard, Adeline Simon, Ricardo Rodriguez de la Vega, Monika Coton, Emmanuel Coton, Marie Foulongne-Oriol, Antoine Branca, Tatiana Giraud

**Affiliations:** Ecologie Systématique Evolution, IDEEV, Bâtiment 680, 12 route RD128, Gif-sur-Yvette, France; Univ Brest, Laboratoire Universitaire de Biodiversité et Ecologie Microbienne, F-29280, Plouzané, France; Laboratoire Interprofessionnel de Production – SAS L.I.P., 34 rue de Salers, Aurillac, France; INRAE, MycSA, Mycologie et Sécurité des Aliments, 33882 Villenave d’Ornon, France; Université Clermont Auvergne, INRAE, Vetagro Sup, UMRF, 20 Côte de Reyne, Aurillac, France; Université Paris-Saclay, INRAE, UR1290 BIOGER, Palaiseau, France; Université Paris-Saclay, CNRS, IRD, UMR Évolution, Génomes, Comportement et Écologie, 91190 Gif-sur-Yvette, France

**Author notes:** These authors jointly supervised the study (Marie Foulongne-Oriol, Antoine Branca, Tatiana Giraud). These authors contributed equally to this work (Thibault Caron, Ewen Crequer). Corresponding author: Ewen Crequer < >; “Tatiana Giraud” < >.

**Keywords:** genetic map, filamentous fungi, translocation, chromosomal rearrangements, breeding, yeast, food, Starship

## Abstract

Elucidating the genomic architecture of quantitative traits is essential for our understanding of adaptation and for breeding in domesticated organisms. *Penicillium roqueforti* is the mold used worldwide for the blue cheese maturation, contributing to flavors through proteolytic and lipolytic activities. The two domesticated cheese populations display very little genetic diversity, but are differentiated and carry opposite mating types. We produced haploid F1 progenies from five crosses, using parents belonging to cheese and non-cheese populations. Analyses of high-quality genome assemblies of the parental strains revealed five large translocations, two having occurred via a circular intermediate. Offspring genotyping with genotype-by-sequencing (GBS) revealed several genomic regions with segregation distortion, possibly linked to degeneration in cheese lineages. We found transgressions for several traits relevant for cheese making, with offspring having more extreme trait values than parental strains. We identified quantitative trait loci (QTLs) for colony color, lipolysis, proteolysis, extrolite production, including mycotoxins, but not for growth rates. Some genomic regions appeared rich in QTLs for both lipid and protein metabolism, and other regions for the production of multiple extrolites, indicating that QTLs have pleiotropic impacts. Some QTLs corresponded to known biosynthetic gene clusters, e.g., for the production of melanin or extrolites. F1 hybrids constitute valuable strains for cheese producers, with new traits and genetic diversity, and allowed identifying target genomic regions for traits important in cheese making, paving the way for strain improvement. The findings further contribute to our understanding of the genetic mechanisms underlying rapid adaptation, revealing convergent adaptation targeting major regulators.

## Introduction

Most traits of agricultural or evolutionary significance are quantitative and genetically complex, involving multiple genes that interact with one another and the environment (Falconer and Mackay 1995). Genomic loci showing diversity associated with quantitative trait variation are called QTLs, for “quantitative trait loci”. Domesticated species are good models for understanding the genomic architecture of quantitative traits involved in adaptation. Indeed, they have been subjected to recent and strong selection on known traits, and display highly contrasting phenotypes compared to wild populations, as well as between different domesticated varieties. In addition to the fundamental interest in understanding the genomic basis of adaptive changes, the agronomically important traits targeted for genetic improvement generally follow quantitative inheritance.

In the QTL mapping approach, genotyping and phenotyping of progenies allow assessing statistical associations between genotypes and phenotypes, thereby allowing identifying genomic regions affecting traits of interest (Falconer and Mackay 1995). The QTL approach has largely advanced our understanding of the genomic architecture of important traits, for example in crops and cattle (Adamczyk et al. 2013; Silva et al. 2014; Kumar et al. 2017). Studies of the genomic architecture of adaptation in crops have revealed that traits involved in domestication were often controlled by only a few QTLs with pleiotropic effects, often being major regulators (Sweeney and McCouch 2007; Baach et al. 2008; Bachlava et al. 2010; Wang et al. 2010; Telias et al. 2011; Wirén and Jensen 2011; Andargie et al. 2014; Johnsson et al. 2014; Wright 2015; Wright et al. 2015; Kongjaimun et al. 2012; Kantar et al. 2017; Somta et al. 2020; Bomblies and Doebley 2006; Martìnez-Ainsworth and Tenaillon 2016). The most emblematic example is the *teosinte branched 1 (tb1)* locus in maize, that determines apical dominance as well as other domestication traits, which is due to a transposable element insertion in a regulatory region (Doebley et al. 1997; Martìnez-Ainsworth and Tenaillon, 2016).

Domesticated fungi used for food production, such as *Saccharomyces cerevisiae* for fermentation, filamentous fungi for cheese maturation or the button mushroom *Agaricus bisporus* for direct human consumption, also represent good models for understanding the genomics of adaptation, in particular as they have small and compact genomes (Gladieux et al. 2014). QTL mapping proved to be an efficient tool for identifying the genes affecting phenotypes impacting technological performances of industrial yeast strains, such as thermotolerance, chemical resistance, dehydration stress tolerance, and phenotypes associated with the fermentation process, such as volatile compounds production, ethanol production, in nitrogen-limited fermentations and in nitrogen consumption and utilization (Nguyen *et al*. 2022; Kessi-Pérez et al. 2020; Eder *et al*. 2018; Wilkening *et al*. 2014; Swinnen, Thevelein and Nevoigt 2012; Liti and Louis 2012; Steinmetz *et al*. 2002). In addition to classic QTLs, traits have also been reported to be impacted in yeasts by aneuploidy (Todd, et al. 2017) and genomic rearrangements (Zimmer *et al*. 2014). QTL mapping has also been successfully used in *A. bisporus,* for identifying the genomic regions controlling color and spore number (Foulongne-Oriol et al. 2012; Imbernon et al. 1996), in the causal agent of the cereal disease *Fusarium* head blight, *Fusarium graminearum*, for identifying the genetic determinants of aggressiveness (Laurent et al. 2021), and in the *Lachancea waltii* yeast, for identifying genomic regions controlling the growth rate in the presence of various drugs (Peltier et al. 2021).

*Penicillium roqueforti* is the mold used for the maturation of all types of blue cheeses worldwide, responsible for the typical blue-veined aspect, and contributing to their specific flavor and aroma, in particular through high levels of proteolytic and lipolytic activities (Moreau 1980; Cerning et al. 1987; Collins et al. 2003). When this study was performed, four populations had been identified in *P. roqueforti,* two populations being used to produce cheeses and two populations being found in molded silage and lumber or in spoiled food (Dumas et al. 2020). The two populations used for cheese maturation each show footprints of bottlenecks and a domestication syndrome, with phenotypes contrasting with those of the non-cheese populations and beneficial for several important aspects of cheese safety, appearance and flavor (Dumas et al. 2020; Caron et al. 2020; Crequer et al. 2023). The two cheese populations each correspond to a clonal lineage, with contrasted phenotypes and diversity levels, and inoculated in different types of cheeses, *i.e.* Roquefort PDO (protected designation of origin) versus other blue cheeses (Gillot et al. 2015; Dumas et al. 2020). The *P. roqueforti* cheese populations produce cheeses with higher percentages of blue area and with higher quantities of desired volatile compounds than non-cheese populations (Caron et al. 2020).

The Roquefort population, found in Roquefort PDO cheeses, displays some level of genetic diversity and exhibits traits beneficial for pre-industrial cheese production, *e.g.* slower growth in cheese models and greater spore production on bread, the traditional multiplication medium (Gillot et al. 2017; Dumas et al. 2020). The Roquefort population also produces higher quantities and more diverse positive aromatic compounds in cheeses, which is due to its faster and different proteolysis and lipolysis activities (Caron et al. 2020; Dumas et al. 2020). In addition, the Roquefort population produced cheese with lower water activity, which could restrict spoiling microorganisms (Caron et al. 2020). Lipolysis and proteolysis are important traits in *P. roqueforti,* involved in cheese maturation, yielding the specific volatile and metabolic compounds responsible for the desired strong and spicy blue cheese flavors (Cerning et al. 1987; Collins et al. 2003; Gillot et al. 2017).

The other cheese population, named non-Roquefort, is constituted by a single clonal lineage, due to a recent strong selection for a single “performant” strain, and is found in all types of blue cheese worldwide except Roquefort PDO cheeses. The non-Roquefort population display phenotypes more suited for industrial cheese production, such as a more efficient cheese cavity colonization ability, higher tolerance to salt, to acidic pH and to lactic acid compared to other populations (Dumas et al. 2020; Ropars et al. 2020; Crequer et al. 2023), and the production of volatiles important for aroma and flavor, *e.g.* methyl ketones (Caron et al. 2020). The non-Roquefort clonal lineage acquired large genomic regions by horizontal transfers (Dumas et al. 2020; Ropars et al. 2015), mediated by giant Starships mobile elements (Gluck-Thaler et al. 2022). These horizontally transferred regions include in particular two very large regions called *Wallaby* and *CheesyTer*, which encompass genes with functions in lactose metabolism or competition against other microorganisms. The non-Roquefort lineage further displays footprints of positive selection in genes involved in volatile compound production (Dumas et al. 2020) and has lost the ability to secrete the mycophenolic acid (MPA) mycotoxin, an immunosuppressant used for preventing transplant organ rejection (Matas et al. 2013), because of a deletion in a key gene (*mpaC*) of its biosynthesis pathway (Gillot et al. 2017; Crequer et al. 2024). *Penicillium roqueforti* can secrete other metabolites, *i.e.* extrolites, including some considered as toxins (*e.g.* Roquefortine C and PR toxin), but not found in blue cheeses, or in such low concentration that they do not cause acute health hazards (Scott, 1981; Fontaine et al. 2015; Hymery et al. 2017). Other *P. roqueforti* extrolites can be of interest, such as the potential antitumor compounds andrastin A and (iso)-fumigaclavine A (Ge et al. 2009; Matsuda et al. 2013). These active compounds may be used by the fungus in the competition against other microorganisms (Jakubczyk et al. 2020; Conrado et al. 2022), and thus avoid cheese spoilage, although little is known on these aspects.

Identifying the genomic regions controlling the phenotypes contrasting between the four *P. roqueforti* populations thriving in different niches would have important fundamental implications for our understanding of the genomic architecture of adaptation and also applied consequences for strain improvement by marker-assisted selection in progenies. However, because of the clonal population structure of the two domesticated cheese lineages, it is not possible to identify the genetic determinants of important traits for cheese making through phenotype/genotype association without producing a recombinant population, which would also allow generating genetic and phenotypic diversity from the two single clonal lineages used in most blue cheeses worldwide (Dumas et al. 2020). Fortunately, sexual reproduction can be induced in *P. roqueforti* (Ropars et al. 2014), the two cheese lineages carry opposite mating types, and some strains have retained some level of sexual fertility, despite the general degeneration in sexual reproduction ability in the cheese populations (Ropars et al. 2016). As the life cycle of *P. roqueforti* has a dominant haploid phase, direct access to this phase allows association genetics to be carried out on first-generation offspring.

Here, we therefore aimed at generating diversity in cheese *P. roqueforti* populations and at identifying the genomic regions controlling the phenotypic differences between the non-Roquefort and Roquefort lineages, and more generally between *P. roqueforti* populations. For this goal, we produced five sexual F1 haploid progenies by crossing six fertile haploid *P. roqueforti* strains from the four originally identified populations at the beginning of the study, focusing particularly on the cross between the two cheese populations, Roquefort and non-Roquefort. We studied in the progenies several phenotypes important for cheese making, *i.e.* lipolysis, proteolysis, growth, color and the production of the main known extrolites: roquefortine C and PR toxin, with its intermediates eremofortins A and B, as well as metabolites with lower toxicity risk, *i.e.* andrastin A, (iso)-fumigaclavine A and MPA (Coton et al. 2020; Chàvez et al. 2023). Depending on the desired cheese product, different values may be wanted for the various traits analyzed. For example, slower growth, lipolysis and proteolysis may allow to store cheeses longer without product degradation, while faster lipolysis may contribute to stronger aromas and faster growth to bluer cheeses (Caron et al. 2020). We analyzed 1,073 offspring with single nucleotide polymorphisms (SNPs) obtained using genotype-by-sequencing (GBS). Because genotyping revealed the presence of the two parental alleles in some genomic regions in the otherwise haploid offspring, we also generated high-quality genome assemblies for the parental strains, to compare the parental genomes and identify genomic translocations. We built genetic maps and ran QTL analyses to identify the genetic architecture of the phenotypes important for cheese making.

## Results

### Five fertile crosses involving at least one cheese strain

We chose the five most fertile crosses among 17 trials involving in each case at least one cheese strain. We identified the isolated spores that were actually recombinant ascospores and not asexual conidia by using 11 microsatellite markers. We focused on the cross between the two cheese populations, *i.e.* Roquefort (R) and non-Roquefort (N), as it may be the most interesting for strain improvement in the cheese making context, and we isolated 387 recombinant offspring (Table 1). The other crosses each involved a cheese strain (either R or N) and a strain from a non-cheese population, either silage (S) or lumber/food spoiler populations (L), and we isolated between 157 and 185 recombinant offspring for each of these four crosses (Table 1).

**Table 1:** The five crosses performed using six strains of *Penicillium roqueforti* from different populations (R for Roquefort, N for non-Roquefort, L for ber/food spoiler, S for silage/food spoiler) with opposite mating types (MAT1-1 and MAT1-2), with the number of isolated and phenotyped offspring and number of marker obtained from GBS genotyping. LCP: « Laboratoire de Cryptogamie, Paris ». *The strain LCP06039 was found to be a mixture of two rphotypes (”dark” and “light”), which was detected after the cross had been performed. ¤These offspring have been genotyped with indels, in addition to S, as all other offspring.

| Cross | MAT1-1 |  | MAT1-2 |  | Ascospore<br>s isolated | Offspring<br>GBS-genotype<br>d | Offspring<br>growth-<br>phenotype<br>d | Offspring<br>color-<br>phenotype<br>d | Offspring<br>lipolysis<br>phenotype<br>d | Offspring<br>proteolysis<br>phenotype<br>d | Offspring<br>phenotyped<br>(mycotoxins<br>) | Offspring<br>used for<br>map<br>construction | Numbe<br>r of<br>marker |
| --- | --- | --- | --- | --- | --- | --- | --- | --- | --- | --- | --- | --- | --- |
|  | Populatio<br>n | Strain | Population | Strain |  |  |  |  |  |  |  |  |  |
| RxN | Roquefort | LCP06136 | non-Roquefort | LCP0613<br>3 | 389 | 384¤ | 307 | 311 | 387 | 382 | 274 | 358 | 2462 |
| RxS | Roquefort | LCP06136 | silage/food<br>spoiler | LCP0604<br>3 | 157 | 157 | 157 | 157 | 157 | 156 | 0 | 148 | 2902 |
| RxL | Roquefort | LCP06136 | lumber/foo<br>d spoiler | LCP0603<br>7 | 185 | 185 | 129 | 131 | 150 | 141 | 120 | 159 | 2943 |
| SxN | silage/food<br>spoiler | LCP06059 | non-Roquefort | LCP0613<br>3 | 176 | 176 | 92 | 92 | 150 | 150 | 75 | 160 | 2860 |
| LxN | lumber/foo<br>d spoiler | LCP06039<br>* | non-Roquefort | LCP0613<br>3 | 171 | 171 | 91 | 91 | 150 | 151 | 0 | 165 | 2618 |

### Transgression and heterosis in phenotype distributions

We measured several phenotypes important for blue cheese production, *i.e.,* mycelium growth, colony color, lipolysis, proteolysis and extrolite production. We analyzed nine variables summarizing these traits in the five progenies, plus seven traits related to extrolite production in three crosses, resulting in 66 trait distributions (9×5+7×3). Among these 66 distributions, 28 did not significantly deviate from normality, 10 best fitted a unimodal distribution and 30 a bimodal distribution (Supplementary Figures 1 and 2; Supplementary Table 1). None of the phenotype distributions were trimodal or amodal (Supplementary Table 1).

The distribution of the five progenies on the principal component analysis based on trait values (Supplementary Figure 3) shows that they represent overall a higher phenotypic diversity than each cross considered separately. A few traits appeared positively or negatively correlated (Figure 1 D), some associations being expected, such as between color traits (*e.g.* blue and red) or between the production of different targeted extrolites (Marcano et al. 2023; Torrent et al. 2017; Rojas-Aedo et al. 2018; Gillot et al. 2017). Other associations were more surprising, such as the negative correlation between proteolysis rate on the one hand, and green and hue level on the other hand (Figure 1 D).

**Figure 1:**
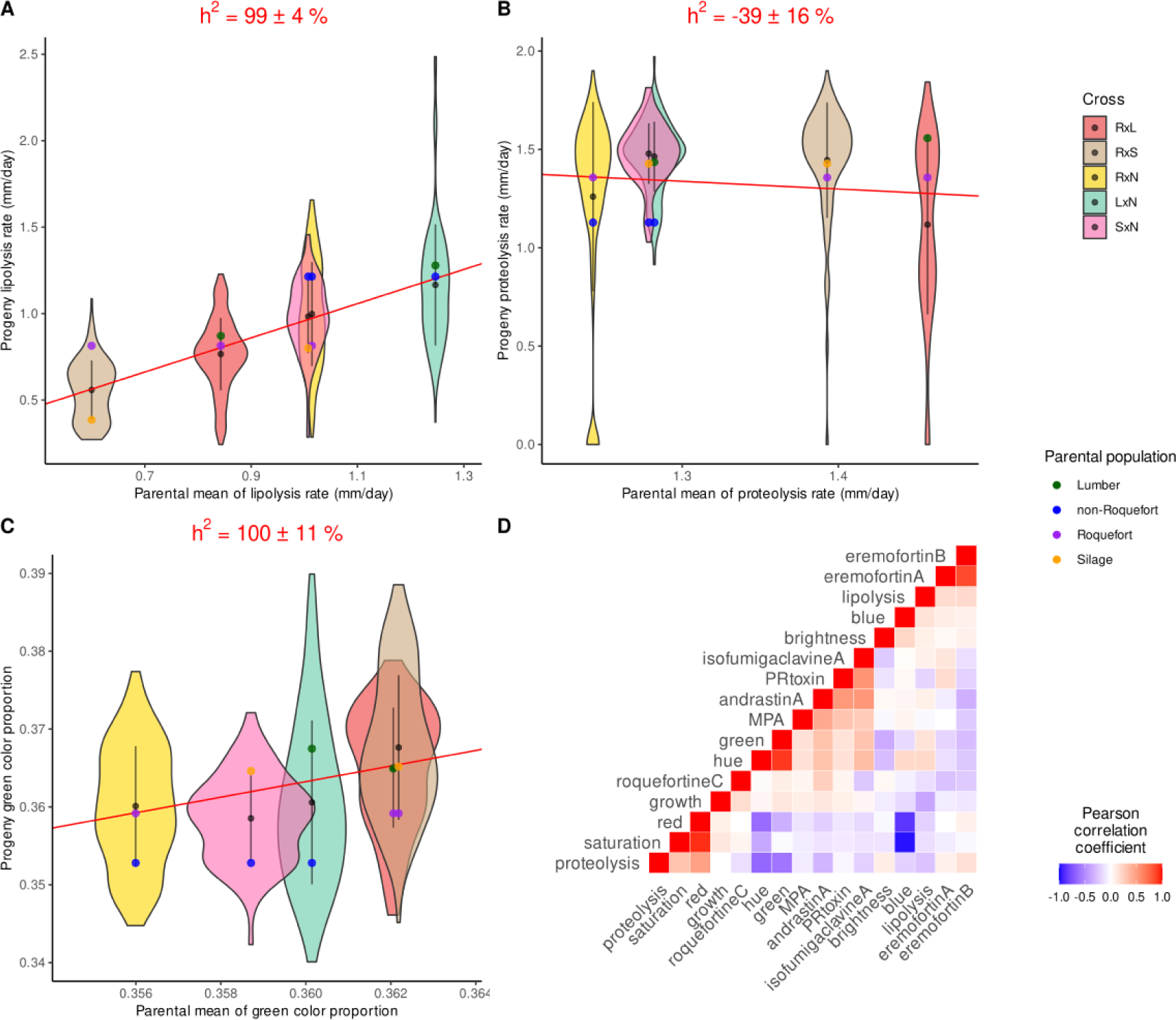
Distributions of phenotypes in the five progenies in *Penicillium roqueforti* as a function of their parental means, for (A) the lipolysis rate (mm/day), (B) the proteolysis rate (mm/day) and (C) the relative green color channel of the progeny; the parental means are plotted along the x axis (RxL in red, RxS in beige, RxN in yellow, LxN in turquoise and SxN in pink). The black point and line represent the mean and standard deviation, respectively, for each progeny. The colored points represent the parental values (lumber in green, non-Roquefort in blue, Roquefort in purple, silage in orange). The red line represents the linear regression across progenies, from which the slope is used to estimate the heritability (h^2^) value given at the top. (D) Pearson correlation coefficients, as a color gradient, between trait values across offspring and crosses.

We found heterosis in the progenies, computed as the difference in mean trait values between the parents and their progenies (Supplementary Table 2). Out of the 66 phenotype distributions, 53 displayed significant positive or negative heterosis. We found negative heterosis in all crosses for the production of PR toxin (the most toxic *P. roqueforti* mycotoxin) and andrastin A, indicating that progenies produced on average less of these two extrolites than their parent mean, which can be beneficial for cheese making. In contrast, we found positive heterosis in all crosses for MPA production, a less toxic extrolite. All tested phenotypes exhibited both positive and negative transgressions across all crosses, *i.e.* with some offspring displaying higher values and others lower values than either parent, which holds promise for the generation of phenotype diversity and strain improvement. We found a large proportion of offspring with low or no PR toxin production in the RxN and SxN crosses. We detected in all crosses negative transgression in roquefortine C production level (*i.e.* the offspring mean was lower than the parental mean) and many offspring with no MPA production, which is promising for the selection of hypotoxinogenic strains.

### High values of phenotype heritability

Narrow-sense heritability (h^2^) represents the part of phenotypic variance explained by additive genetic variance, *i.e.* the part on which selection can act. The narrow-sense heritability of growth rate, lipolysis, proteolysis and color across the five progenies, estimated by regressions of the progeny trait means on the parent trait means, were relatively high, with for example two-thirds of traits with h² above 30% (Table 3). Some estimates were below zero or higher than 100% (Figure 1), but narrow-sense heritability estimates based on regressions between offspring and parent trait means may not be highly reliable with only five crosses, and even only three crosses in the case of the targeted extrolites. For extrolite production, we also estimated broad-sense heritability (H²) as we measured two replicates for each strain (Supplementary Table 3). The broad sense heritability estimates for extrolite production were high, ranging between 91.3 % for roquefortine C and 99.6% for isofumigaclavine A, which is promising for selection (Supplementary Table 3).

### Translocations between parental genomes

The comparison of high-quality genome assemblies revealed five large genomic chromosomal rearrangements between the parents in four crosses (LxN, RxN, RxS, RxL; Figure 2), which had impacts on genetic maps. These translocations ranged from 60 to 530 kb in size. The two silage parents and the non-Roquefort parent showed syntenic genomes between each other, suggesting that they kept the ancestral arrangement order. We could therefore infer that four of the translocations (L1, L2, L3 and L4) occurred in the lumber population while the fifth one, R1, occurred in the Roquefort population (Figure 2). We detected no particular transposable element family or clusters at the margin of the translocations or DUF domains specific to Starship elements.

**Figure 2:**
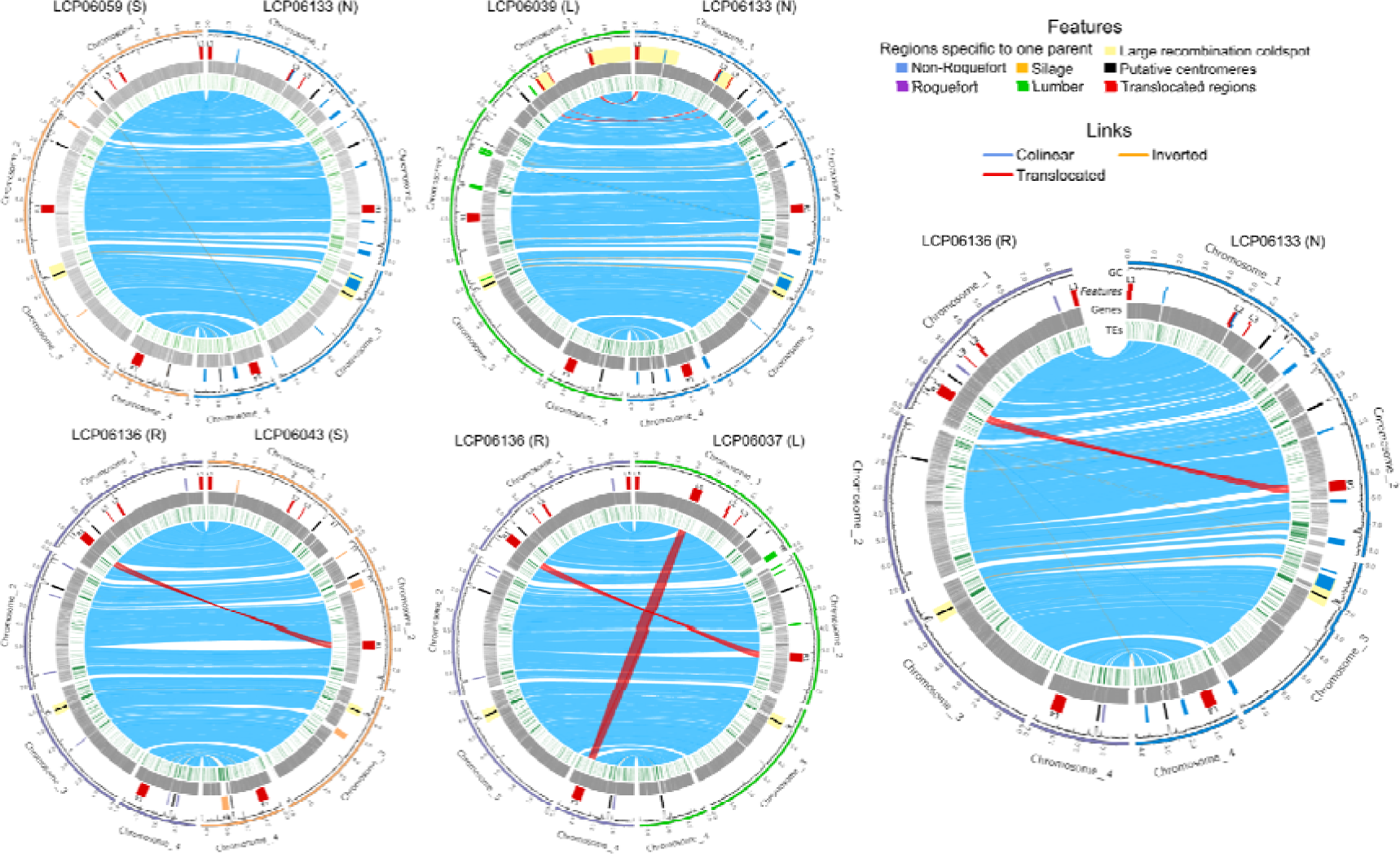
Genome comparisons between parental strains for each cross. Chromosomes are represented with colors corresponding to the population-of-origin of the strains: light orange, blue, purple and green for silage (S), non-Roquefort, (N) Roquefort (R) and lumber (L) populations, respectively. From outside to inside, the tracks represent the i) GC content, ii) translocated regions of more than 50kb between parental genomes in red (with their IDs, the first letter indicating the population-of-origin of the parent in which the translocation occurred), specific regions that are lacking in other genomes (*i.e.* acquired horizontally transferred regions) with their color indicating the populations in which they are present, putative centromeres in black, and large recombination cold spots in light yellow, iii) predicted genes in grey and iv) transposable elements in green. Orthologs between genomes are linked one to each other, with blue, orange and red links, indicating synteny, inversions or >50kb translocations, respectively.

The L1 and L4 regions displayed internal rearrangements between their copies in different genomic locations: the last 3’ portion of each of the L1 and L4 regions in the parents with the ancestral arrangement was located in 5’ of the L1 and L4 regions in the translocated parent (Supplementary Figure 4). This internal rearrangement suggests that the translocation occurred via a circular intermediate mechanism, with an insertion cutting site in each of the L1 and L4 regions different from their excision site (Supplementary Figure 4).

Both parental alleles for the R1 region insertion were found together in 38, 31 and 24 % of the RxN, RxL and RxS offspring in GBS data, respectively (Figure 2). The Oxford Nanopore long-read mapping depths confirmed the presence of two R1 region copies in the six analyzed offspring from the RxN cross with both R1 region parental alleles detected in GBS data. Similarly, the long-read mapping confirmed that the five offspring with a single parental allele in GBS data carried a single copy of the translocated region in their genomes. The absence of this translocated region is likely lethal as we did not detect any offspring with no R1 copy, while some recombination events should have generated offspring with none of the two copies. We detected no offspring with both parental alleles nor an absence of the region for the translocations L1 to L4, indicating that both the duplication or absence of these regions lead to inviability.

We checked that the presence of the two R1 copies in haploid genomes was stable after culture and replication of 24 offspring of the RxN cross carrying the two parental alleles. After cultivating these 24 offspring from the RxN cross for one week on plates and transferring conidia from the edge of the colony to new plates 19 times, PCRs showed that the cultivated lineages still carried the two R1 parental alleles. We did not detect any RIP footprints (C-to-T repeat-induced point mutations) in the R1 regions of the six offspring with a duplicated R1 region and with high-quality genome assemblies. However, RIP footprints would only be expected in the following generation (F2), after a sex event involving genomes with the duplicated region, as RIP is known to occur during the short dikaryotic phase of sexual reproduction in ascomycetes (Galagan and Selker 2004).

### Genetic maps, recombination pattern and segregation biases

We identified between 2,462 and 2,943 reliable markers in the progenies across the five crosses, evenly distributed along the non-Roquefort reference genome. For constructing genetic maps, we excluded redundant markers, *i.e.* those physically close and with identical segregation patterns in a given cross. We thus constructed the genetic maps with 1,448 markers for the RxN cross, and 676 to 785 markers for the other crosses with fewer analyzed offspring (Table 2). The genetic maps for the five crosses displayed lengths between 775 and 913 cM, with a mean of 857 cM (Table 2). We detected four linkage groups in each genetic map, corresponding to the four chromosomes in the parental genome assemblies (Figure 3). The mean number of crossing-overs per chromosome was 2.15, with a maximum of 2.58 for chromosome 1, the longest chromosome, and a minimum of 1.25 for chromosome 4, the smallest chromosome. The mean recombination rate was estimated between 27.0 cM/Mb and 31.8 cM/Mb, with a mean of 29.8 ± 1.8 cM/Mb (Table 2). The LxN cross had a particularly low mean recombination rate, mainly due to the presence of two cold spots of recombination on chromosome 1 (indicated by a plateau in the plot of genetic map positions against genomic positions; empty rectangles in Figure 3), with local recombination rates of 6.7 cM/Mb and 3.1 cM/Mb, respectively (Table 2). The edges of the two cold spots corresponded to the translocated regions in the chromosome 1 of the LCP06039 strain (regions L1, L2, L3, Figure 2 and 3). This is likely due to the inviability of offspring carrying either zero or two copies of the translocated region because of recombination events between their insertion sites (Figure 2). Other regions with low or no recombination rates corresponded to horizontally transferred regions only present in the reference genome, putative centromeres, other translocated regions and TE-rich regions (Figure 2 and 3).

**Table 2:** The five cros 1500 ses performed using six strains of *Penicillium roqueforti* from different populations (R for Roquefort, N for non-Roquefort, L for lumber/food spoiler, S for silage/food spoiler), the linkage group and the total, the number of markers used for genetic map construction, the total length of the linkage group, the average and maximum spacing in cM between markers, the mean recombination rate in cM/Mb, and the reference chromosome and genome size in Mb.

| Cross | Linkage group | Number of markers | Mean number of crossing over | Length (cM) | Average spacing (cM) | Maximum spacing (cM) | Mean recombination rate (cM/Mb) | Reference size (Mb) |
| --- | --- | --- | --- | --- | --- | --- | --- | --- |
| **SxN** | L.1 | 213 | 2.74 | 274 | 1.3 | 8.8 | 32.21 | 8.51 |
|  | L.2 | 242 | 2.49 | 249 | 1 | 12.8 | 29.97 | 8.31 |
|  | L.3 | 189 | 1.97 | 197 | 1 | 8.8 | 25.51 | 7.72 |
|  | L.4 | 94 | 1.19 | 120 | 1.3 | 8.8 | 28.47 | 4.22 |
|  | overall | 738 | 8.39 | 841 | 1.1 | 12.8 | 29.25 | 28.75 |
| **LxN** | L.1 | 161 | 1.98 | 199 | 1.2 | 9.2 | 23.39 | 8.51 |
|  | L.2 | 248 | 2.55 | 255 | 1 | 9.2 | 30.69 | 8.31 |
|  | L.3 | 164 | 1.95 | 195 | 1.2 | 6.7 | 25.25 | 7.72 |
|  | L.4 | 103 | 1.28 | 128 | 1.3 | 8.6 | 30.36 | 4.22 |
|  | overall | 676 | 7.75 | 776 | 1.2 | 9.2 | 26.99 | 28.75 |
| **RxN** | L.1 | 466 | 2.78 | 279 | 0.6 | 7.3 | 32.79 | 8.51 |
|  | L.2 | 435 | 2.48 | 248 | 0.6 | 8.7 | 29.85 | 8.31 |
|  | L.3 | 378 | 2.22 | 223 | 0.6 | 9.6 | 28.88 | 7.72 |
|  | L.4 | 179 | 1.25 | 126 | 0.7 | 8.7 | 29.89 | 4.22 |
|  | overall | 1458 | 8.74 | 875 | 0.6 | 9.6 | 30.43 | 28.75 |
| **RxS** | L.1 | 191 | 2.74 | 275 | 1.4 | 11.7 | 32.32 | 8.51 |
|  | L.2 | 242 | 2.47 | 248 | 1 | 13.9 | 29.85 | 8.31 |
|  | L.3 | 227 | 2.55 | 256 | 1.1 | 12.4 | 33.15 | 7.72 |
|  | L.4 | 115 | 1.33 | 133 | 1.2 | 7.5 | 31.55 | 4.22 |
|  | overall | 775 | 9.10 | 913 | 1.2 | 13.9 | 31.75 | 28.75 |
| **RxL** | L.1 | 186 | 2.64 | 264 | 1.4 | 14.9 | 31.03 | 8.51 |
|  | L.2 | 295 | 2.66 | 267 | 0.9 | 13.5 | 32.14 | 8.31 |
|  | L.3 | 181 | 2.30 | 231 | 1.3 | 9.5 | 29.91 | 7.72 |
|  | L.4 | 129 | 1.19 | 119 | 0.9 | 12.9 | 28.23 | 4.22 |
|  | overall | 791 | 8.79 | 881 | 1.1 | 14.9 | 30.64 | 28.75 |
| **Crosses mean** | L.1 | 243.4 | 2.58 | 258.2 | 1.18 | 10.38 | 30.348 | 8.51 |
|  | L.2 | 292.4 | 2.53 | 253.4 | 0.9 | 11.62 | 30.5 | 8.31 |
|  | L.3 | 227.8 | 2.20 | 220.4 | 1.04 | 9.4 | 28.54 | 7.72 |
|  | L.4 | 124 | 1.25 | 125.2 | 1.08 | 9.3 | 29.7 | 4.22 |
|  | overall | 887.6 | 8.55 | 857.2 | 1.04 | 12.08 | 29.812 | 28.75 |

**Figure 3:**
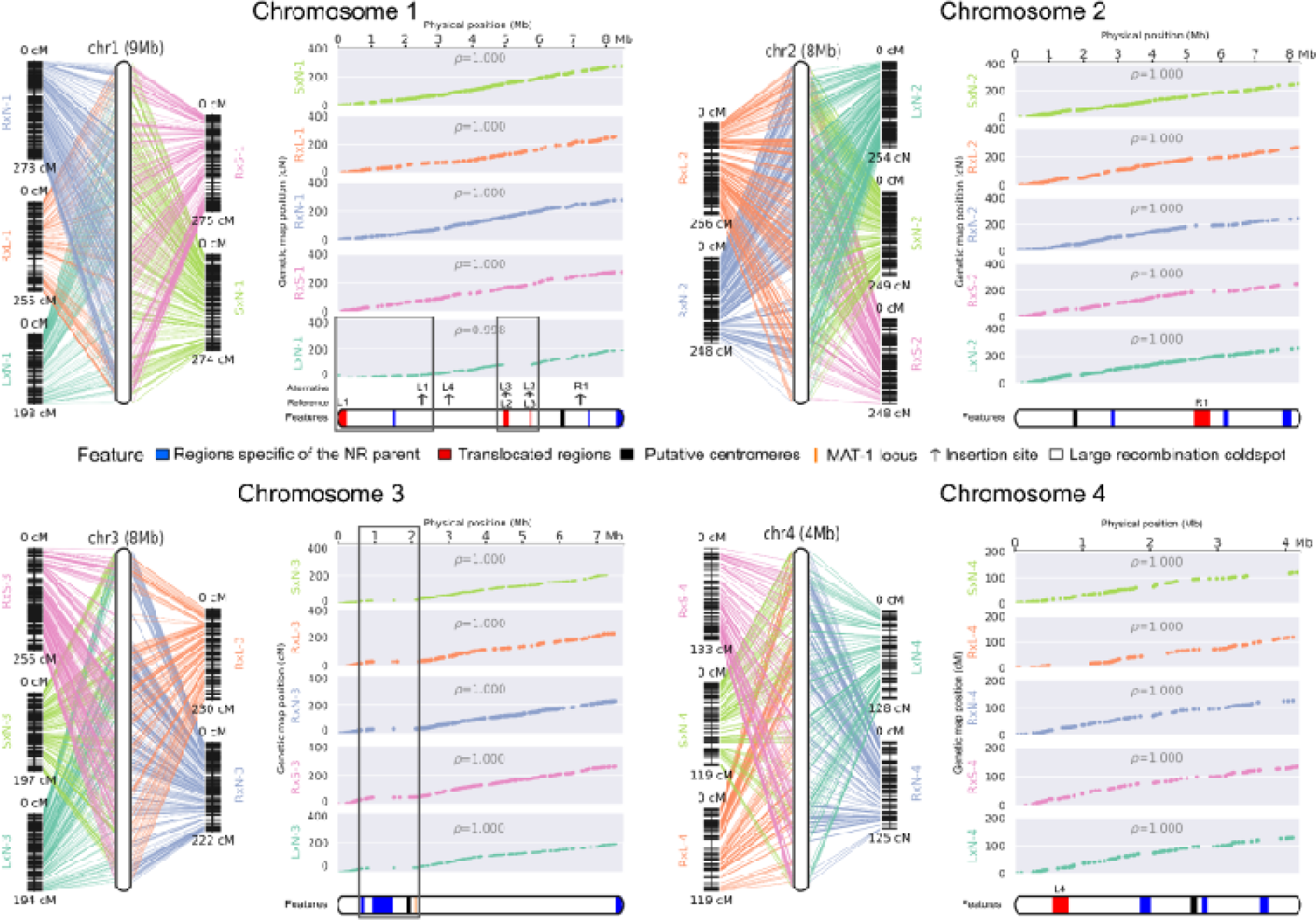
Genetic and physical maps for the four chromosomes of *Penicillium roqueforti,* obtained by analysing five inter-population crosses. Each panel corresponds to an individual chromosome, line graphs on the left represent linkage groups (one color per cross, with cross IDs as in Table 1; note that different parents were used for the silage and lumber populations between crosses); the lines link markers to chromosomes of the reference genome (LCP06133 non-Roquefort parent). In each panel, plots on the right represent the relationship between genetic and physical distance, with one plot per cross and the same color code as in left panels. At the bottom of each panel, the reference chromosome is represented, with filled black, red and blue rectangles representing the putative centromeres, the translocated regions and the horizontally transferred regions of more than 50kb when present, respectively. The red translocated regions are labelled as in Figure 2. The orange bar in chromosome 3 represents the mating-type locus position. The empty black rectangles indicate large recombination cold spots situated between translocated or parent-specific regions.

In the five crosses, large regions presented segregation biases, with under-representation of alleles from the cheese populations (*i.e*. either from the Roquefort or non-Roquefort parents; Figure 4). Such segregation biases could allow the purge of deleterious mutations in the cheese clonal lineages, but could also render the selection of valuable alleles that are in linkage with deleterious alleles in these regions more challenging. The segregation bias against the non-Roquefort parent allele in the second half of chromosome 4 (between 3.5Mb and 4.2Mb) was present in all crosses involving the non-Roquefort parent (Figure 4) and was very strong (>90% of the alternative allele in all three crosses). Such strong under-representation of alleles from the cheese population in all of these three crosses can be due to the lower fertility of domesticated populations (Ropars et al. 2016), with likely deleterious alleles accumulated through clonal replication that lead to low viability of the ascospores carrying them. We observed another segregation bias with strong over-representation (> 90%) of silage parental alleles, in the RxS and SxN crosses, in chromosome 1 from 4.2 to 4.5Mb (Figure 4). The under-represented allele, that of the cheese parents, displayed no segregation bias in the other crosses involving the cheese parents (Figure 4), suggesting the presence of either a selfish element (*e.g.* a spore killer) or a highly beneficial allele for early growth in the corresponding region in the silage/spoiled food parents rather than a deleterious one in the cheese parent. This interpretation is supported by the over-representation of the silage allele in two different crosses and the lack of bias in the RxL and LxN crosses in this region, while these observations would not be expected if the cheese parents had deleterious alleles in this region. The segregation bias with over-representation of Roquefort alleles at the end of chromosome 1 is likely due to the lack of the R1 translocated region that leads to offspring inviability. Symmetrically, the alleles from the other parent tended to be over-represented at the R1 locus on chromosome 2, but not significantly so. This suggests that the location of R1 may be more advantageous on chromosome 1 than on chromosome 2.

**Figure 4:**
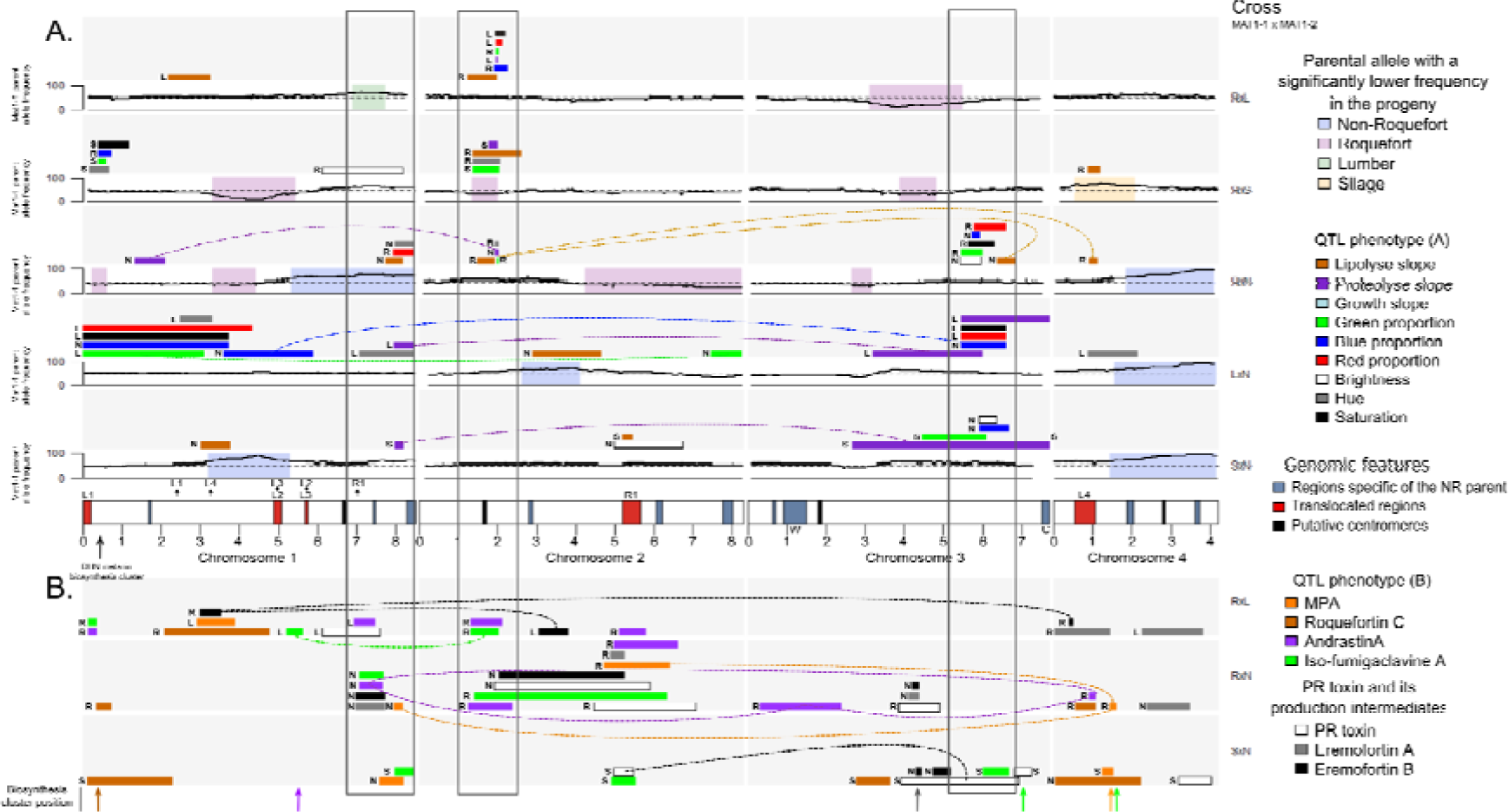
Representation of identified quantitative trait loci (QTLs) and segregation distortion along the four chromosomes of *Penicillium roqueforti* in five progenies. Only QTLs explaining more than 5% of the total variance are represented. The color of the bars indicates the considered trait while the letter indicates the parent having the alleles increasing the trait value. Dotted lines link the QTLs with significant interactions and color indicates the phenotype. The cross ID is indicated on the right, the first parent carrying the MAT1-1 mating type. (A) QTLs identified for phenotypes linked to lipolysis, proteolysis, growth and colony color for the five crosses (in lines, indicated on the right with the same code as in Table 1, note that different parents were used for the silage and lumber populations between crosses). Black curves represent the proportion in progenies of the allele from the parent carrying the MAT1-1 mating type. Transparent colored rectangles on curves indicate regions with significant segregation distortion, their color corresponding to the under-represented parental allele: blue, purple, green and yellow for non-Roquefort, Roquefort, lumber/spoiled food and silage/spoiled food parents, respectively. The x-axis represents genomic physical positions. At the bottom, the four chromosomes of the reference genome (LCP06133) are represented, with rectangles representing genomic features: large horizontally transferred regions specific to the reference genome in blue, and translocated regions in the reference genome in red (with same IDs as in Figure 2). The location of the horizontally transferred regions (W for *Wallaby* and C for *CheesyTer*) are indicated. The location of the dihydroxynaphthalene (DHN) melanin production cluster is indicated by a black arrow. The large vertical empty black rectangles indicate pleiotropic QTL regions, with effects on multiple traits. (B) QTLs identified for extrolite production (with colors corresponding to the different toxins), analyzed in three crosses (SxN, RxN and LxN). The locations of the gene clusters of the toxin biosynthesis pathways are indicated by arrows with the same color code (the PR toxin cluster being in gray, eremofortin A and B being intermediates).

### Detection of QTL with pleiotropic effects

We investigated QTLs for the nine traits associated with lipolysis, proteolysis, growth and color in all descendants, as well as for the seven traits associated with extrolite production specifically in the RxN, RxL and SxN crosses (Supplementary Table 4). We found QTLs for all tested phenotypes except for growth rate (Figure 4; Supplementary Table 4), indicating that marker-assisted selection for these traits may be performed for strain improvement and diversification. We detected 123 QTLs across all phenotypes and crosses, 109 explaining more than 5% of the variance (hereafter called “major QTLs”). We identified 58 major QTLs for the eight parameters related to lipolysis, proteolysis and color, across the five crosses, with 8 to 17 QTLs per cross, with a mean of 1.3 major QTLs and a maximum of 3 major QTLs per trait and cross. For extrolite production, we identified 13 to 23 major QTLs across the three progenies analyzed for these traits, with a mean of 2.4 and a maximum of 5 major QTLs per extrolite and cross (Supplementary Table 4, Figure 4).

The identified QTLs and the direction of their effects were in agreement with the past occurrence of selection for color, proteolysis, lipolysis and extrolite production in cheese populations. We indeed identified QTLs impacting colony color, bluer colonies being associated with cheese parent alleles, and in particular in the genomic region containing the dihydroxynaphthalene-melanin biosynthesis cluster, known to be involved in melanin production (Figure 4). The cheese parent alleles were also both associated with slower proteolysis and faster lipolysis, in connection with firmer texture and longer cheese storage and the production of typical blue cheese flavors, respectively. We identified QTLs for the production of three extrolites, MPA, PR toxin and roquefortine C, the non-Roquefort alleles being associated with lower production levels (Figure 4). This is consistent with a selection for bluer color, more efficient lipolysis and less efficient proteolysis in domesticated lineages, as well as lower extrolite production levels.

Most of the major QTL regions presented pleiotropic effects. The 109 major QTLs were indeed not distributed homogeneously across the genome, with instead regions carrying clustered QTLs for different phenotypes (Figure 4, Supplementary Figure 5). We considered as pleiotropic the regions presenting overlaps in QTL regions of at least two different trait classes in a given cross (proteolysis, lipolysis, color and extrolite production). We identified five such regions, among which three regions displayed pleiotropic impact on the four trait classes, *i.e.* proteolysis, lipolysis, color and extrolite production, across the different progenies, and also within some progenies for multiple traits (empty vertical rectangles in Figure 4, Supplementary Figure 5). For the pleiotropic region on chromosome 2, the Roquefort allele was associated with faster lipolysis and lower proteolysis. This QTL region displayed significant interactions for lipolysis with two other QTLs, in chromosomes 3 and 4; such interactions between loci in determining a trait value constitute positive epistasis. The same region also impacted color. For the pleiotropic regions detected in chromosomes 1 and 3, the non-Roquefort alleles were associated with slower proteolysis, faster lipolysis and bluer colony color (Supplementary Table 4, Figure 4). We further found epistasis for other types of phenotypic traits, particularly between different QTLs controlling proteolysis (Supplementary Table 4, Figure 4). QTL interactions had the same sign (positive or negative) across the various crosses in some cases and different signs in other cases, indicating that epistatic interactions are dependent on the genetic background. Such epistasis could impair selection, as recombination breaks up allelic combinations, while specific allelic combinations are beneficial under positive epistasis.

We also identified QTLs for extrolite level production, including some outside of their known biosynthesis gene clusters, suggesting that these QTLs correspond to *trans*-acting regulators. For three extrolites (MPA, PR toxin and roquefortine C), we identified QTLs with intervals included in their biosynthesis cluster, with non-Roquefort alleles associated with lower production levels. The lower MPA production level associated with the non-Roquefort allele is likely due to the deletion in the *mpaC* gene that was identified in the non-Roquefort population (Gillot et al. 2017; Crequer et al. 2024). We also found a QTL in a region including the PR toxin biosynthesis gene cluster, with lower production level of the PR toxin but accumulation of its production intermediates, eremofortins A and B, associated with the non-Roquefort allele at the PR toxin biosynthesis cluster. This suggests a disruption in the cluster driving low production levels of the toxin and accumulation of its intermediates in offspring presenting the non-Roquefort allele likely due to a premature STOP codon in the ORF11 gene of the cluster (Crequer *et al*. 2024). The offspring harboring a non-Roquefort allele for the various biosynthesis gene clusters displayed lower production levels of the two most toxic *P. roqueforti* extrolites*, i.e.* PR toxin and roquefortine C, and of the less toxic MPA extrolite, which is consistent with a selection for an hypotoxinogenicity trait in the non-Roquefort population (Fontaine *et al*. 2015; Hymery *et al*. 2017; Crequer *et al*. 2024). Most of the QTLs associated with extrolite production were not located within their biosynthesis gene cluster, which suggests the implication of *trans*-acting regulators. These QTLs were often shared between different extrolites, suggesting a co-regulation of their pathways, which could facilitate the selection for low extrolite production in strains bred for cheese production. In particular, isofumigaclavine A production shared five of its eight major QTLs with andrastin A production, suggesting co-regulation of the production of the two extrolites. We detected QTLs for all tested extrolites, except roquefortine C, in the R1 region for its two insertion sites (Figure 4), with lower production associated with the presence of the R1 region. Such a QTL clustering suggests the presence of a master regulator of extrolite production in the R1 region. We in fact found an ortholog of a master regulator gene in the R1 region, namely the *srk1* gene, regulating osmotic and oxidative responses in fungi (Marquina *et al*. 2022). The selection of offspring with two R1 regions could thus allow further decreases in overall extrolite content, and in particular that of the studied mycotoxins in cheese.

We detected an andrastin A QTL next to the *Wallaby* horizontally transferred region (HTR) in the RxN cross and a proteolysis QTL in the region containing the *CheesyTer* HTR in each of the SxN and LxN crosses (Figure 4). Multiple other phenotypes have previously been suggested to be controlled by these HTRs, such as milk sugar metabolism and interactions with other microorganisms (Ropars *et al*. 2015), but the indel nature of these HTRs may render hard to detect their associations with phenotypes, as markers are lacking in one of the parents.

We also checked whether QTL regions encompassed other regulators known to be involved in extrolite production (*prlaeA,* Marcano *et al*. 2023, *pcz1,* Gil-Durán *et al*. 2015, and *sfk1*, Torrent *et al*. 2017) as well as other essential phenotypes in *P. roqueforti*, such as conidiation or growth rate. We located *pcz1* in the pleiotropic region of chromosome 2, encompassing QTLs for andrastin A production and color, melanin being involved in both color and conidiation (Gil-Durán *et al*. 2015); *pcz1* could therefore be a candidate for the QTLs controlling andrastin A production and color via expression regulation. While no QTL interval overlapped with *prlaeA*, a QTL region controlling MPA production in one cross encompassed *sfk1*. However, this QTL region overlaps with a shorter MPA QTL present in another cross without *sfk1* in the confidence interval, which suggests no implication of *sfk1* in MPA production regulation.

The QTL intervals were too large to identify particular genes or functions beyond the horizontal gene transfers, biosynthesis gene clusters of the targeted extrolites or other *a priori* candidate genes. Indeed, with on average one gene every 3kb in the annotated Roquefort parental strain and a median QTL length of 760 kb, half of the QTL regions are expected to encompass more than 250 genes. We therefore only tried to identify candidate genes in the smallest QTL intervals explaining more than 20% of the phenotypic variance. We found an enrichment in genes involved in carbohydrate catabolism for a QTL associated with color in the pleiotropic region of chromosome 2 (g3621 to g3634), and a pepsin gene (endopeptidase, g5778) for a QTL associated with proteolysis in chromosome 3. Carbohydrate catabolism and pepsin are of paramount importance for secondary starters, to use the remaining lactose residues to establish themselves early in the cheese matrix (Cerning *et al*. 1987), and to degrade caseins and therefore to participate in proteolysis (Grippon *et al*. 1977), respectively.

## Discussion

In this study, we aimed at generating genetic and phenotypic diversity in the blue cheese fungus *P. roqueforti* by analyzing haploid F1 progenies from five crosses between cheese or non-cheese strains, and identifying QTLs for key traits in cheese making. This will be essential for both strain breeding purposes and for studying the genomic architecture of traits selected during domestication. Generating diversity and reshuffling alleles is crucial as deleterious mutations have accumulated in the two clonal lineages currently used for cheesemaking. Moreover, their low diversity raises concern for the sustainable use of *P. roqueforti* for cheese making. We successfully generated offspring by crossing strains from different populations of *P. roqueforti*, despite the known low fertility of the cheese populations (Ropars *et al*. 2016). The progenies displayed a broad diversity for the phenotypes important for cheese making, in terms of growth, color, proteolysis and lipolysis rates, as well as extrolite production, with more extreme trait values than the parents. We detected QTLs for all these traits except growth, often displaying pleiotropic effects. These findings will be highly valuable for strain breeding, and further suggests that humans have domesticated this cheese fungus by selecting major regulators, as reported in crops (Telias *et al*. 2011; Wirén & Jensen, 2011; Wang *et al*. 2010; Sweeney & McCouch, 2007; Wright, 2015; Wright *et al*. 2015; Baach *et al*. 2008; Johnsson *et al*. 2014; Bachlava *et al*. 2010; Andargie *et al*. 2014; Kongjaimun *et al*. 2012; Kantar *et al*. 2017; Somta *et al*. 2020; Bomblies & Doebley, 2006; Martìnez-Ainsworth & Tenaillon, 2016).

### Variability in phenotypes in progenies, with heterosis and positive transgression for most traits

We found large phenotypic variation, transgression and heterosis in progenies for multiple relevant phenotypes for blue cheese production, *i.e*. mycelium growth, lipolysis, proteolysis and extrolite production. Progenies displayed transgression, with the trait values of some offspring being much more extreme than those of the parents. All progenies also exhibited heterosis, indicating mean trait values in offspring that may be better than the mean value of the parents. Both transgression and heterosis are promising for strain improvement and diversification. Furthermore, we detected QTLs for most traits, indicating that selection is possible, and even marker-assisted selection. The generation of variation is particularly important in *P. roqueforti*, as the two main cheese populations, Roquefort and non-Roquefort, have lost most of their genetic diversities due to recent bottlenecks followed by strong selection in industrialization times (Dumas *et al*. 2020). These clonal lineages even degenerate in terms of sexual fertility and probably for other traits of interest for cheese making (Ropars *et al*. 2016). Sexual reproduction is the only way to purge deleterious mutations that accumulate during clonal propagation. The progeny between the Roquefort and non-Roquefort strains is particularly promising, as a European regulation requires a food safety assessment (“Novel food” law; UE No 2015/2283) for the use of microorganisms for food production except if they have already been used for food before May 1997. The cross of parental strains already used for cheese production for decades should therefore meet the European Union requirements. The other crosses may also be of great interest, even for the European market, as the regulation might be interpreted in terms of species rather than strains, as already done for example in wine and beer yeasts.

### Translocations and segregation biases

The obtained high-quality genome assemblies led to the identification of five translocations between the parental strains, having occurred independently in three lineages. We also identified offspring carrying twice the R1 rearranged region. For these translocations, we did not find any offspring completely lacking them, indicating that their absence in genomes may lead to offspring unviability. For the L1, L2, L3, and L4 translocations, we detected no offspring with twice the regions either, implying that their duplications are also lethal. We detected lower production levels for the targeted metabolites in offspring carrying two copies of the R1 region. These offspring may therefore be highly valuable for cheese making, especially as the presence and locations of the translocations were stable through multiple replications and culture events. For further improvement, it should however be checked that future crosses with these offspring will not induce RIP in the duplicated region. RIP footprints (*i.e*. mutations from CA to TA in repeated sequences) have indeed been already detected in *P. roqueforti* (Cheeseman & Ropars *et al*. 2014).

We detected no particular transposable element types or abundance at the margin of the translocations. In particular, we detected no DUF3435 domain, typical of Starship elements, that move within and between fungal genomes via a circular intermediate (Bucknell & MDonald, 2023) and are involved in horizontal gene transfers in *P. roqueforti* (Gluck-Thaler et al. 2022). Nevertheless, the L1 and L4 translocations appeared to have occurred via a circular form. These regions may therefore correspond to a new type of large mobile element with a circular intermediate. These findings reinforce the view that fungal genomes are highly dynamic, with frequent gene movements, within genomes, between individuals or even between species, and that this contributes to adaptation (Bucknell & MDonald, 2023).

For several translocated regions, either their lack or duplication in genomes led to to unviability, which induced the cosegregation of the two insertion sites, potentially generating linkage drag in future breeding, *i.e.* cosegregation of desired and undesired alleles in progenies. For other translocated regions, only their absence was lethal, and this led to segregation biases in progenies. We detected additional segregation biases, in other genomic regions, mostly with under-representation of the parental strain from the parental domesticated cheese populations. Such biases are likely due to degeneration in the cheese clonal lineages, with alleles less fit than in the wild populations, especially under sexual reproduction and ascospore germination (Ropars *et al*. 2016). We detected a strong segregation distortion in chromosome 4, with an absence of the non-Roquefort alleles in the progeny. This suggests an unviability of the ascospores carrying the non-Roquefort allele, in agreement with the low fertility of the non-Roquefort population, showing high post-mating sterility, *i.e.* formation of cleistothecia without ascospores (Ropars *et al*. 2016). The region with segregation distortion in chromosome 4 contained two orthologs of genes involved in sporulation in the *Saccharomyces cerevisiae* yeast, g9043 and g9061 coding for SPS22-like (Coluccio *et al*. 2004) and RMD1-like proteins (inferred by electronic annotation, UniprotKB-KW; Enyenihi & Saunders, 2003), suggesting that the region contains genes essential for sexual reproduction. The region with segregation distortion in chromosome 4 also included genes with premature stop codons in the non-Roquefort parent, although with either no predicted function or a function with *a priori* no relationship with sexual reproduction (g8973 and g8998). In the case of the RxS and SxN segregation bias in chromosome 1, the under-represented Roquefort and non-Roquefort alleles displayed no segregation bias in the other crosses involving the same Roquefort and non-Roquefort parents. This indicates that the bias depends on the genetic background and may therefore be due to epistatic interactions leading to a deleterious effect or a selfish meiotic drive element (Grognet *et al*. 2014; Turner & Perkins, 1979).

### QTL identification and the genomic architecture of adaptation

We identified QTLs for all phenotypes but growth, confirming that selection of these traits for strain improvement can be considered. There were only a few major QTLs per trait and cross, which should facilitate strain breeding. The QTLs for color, lipolysis and proteolysis on the one hand, and for extrolite production on the other hand, clustered in the same genomic regions, suggesting pleiotropy, or clustering of genes with effects on these different pathways. The three pleiotropic regions affecting color, and lipid and protein degradation have the same effects, with faster lipolysis, slower proteolysis and bluer colony color for the cheese strain allele. This is consistent with a selection for a bluer color, stronger flavors and a longer conservation in domesticated lineages of *P. roqueforti* for cheese making. Furthermore, such pleiotropy or gene clustering may render selection for these traits easier or more challenging depending on whether alleles coding for desired and unwanted traits are associated. Actually, pleiotropic regions controlling multiple important traits for cheese making may be the result of human selection targeting major regulators, and their finding informs on the genomic architecture of adaptation. The control of traits involved in domestication by only a few QTLs and their pleiotropy have been reported in animals and plants, often due to major transcription regulators, human selection having targeted “masterminds” (Martìnez-Ainsworth & Tenaillon, 2016), but also to the genetic determinant of important crop traits being clustered in genomes (Wright, 2015; Kantar *et al*. 2017). Here, we show that such pleiotropic regions are present in two independently domesticated lineages of *P. roqueforti,* with the same type of effect on the phenotype, suggesting evolutionary convergence (Ropars & Giraud, 2022) and selection of major regulators with pleiotropic effects. Known shared regulators of various secondary metabolites include *prlaeA*, *sfk1* and *pcz1*, their silencing or knock-out decreasing the production of roquefortine C, andrastin A et MPA (Marcano *et al*. 2023; Torrent *et al*. 2017; Rojas-Aedo *et al*. 2018). Over-expression of *pcz1* decreases the production of roquefortine C and andrastin A, and increases the production of MPA; *pcz1* also impacts growth, conidiation and spore germination (Gil-Durán *et al*. 2015); *pcz1* was found in one of the pleiotropic regions with an impact on andrastin A production and on color, which is mostly due to conidia pigmentation.

In addition, we found QTL regions containing the biosynthesis gene clusters involved in PR toxin, MPA and roquefortine C production. In the RxS and LxN crosses, we found a QTL affecting the colony color in a region containing the DHN melanin biosynthesis gene cluster, known to code for the production of *P. roqueforti* conidia pigment (Cleere *et al*. 2024; Seekles *et al*. 2021). Besides these *a priori* candidates, the QTL analysis did not point to particular genes or regulatory regions, as the regions with significant impacts were large, encompassing many genes, which impaired searching for relevant gene functions. Furthermore, as shown in plant domestication, several important traits have evolved due to transposable element insertions in regulatory regions (Martìnez-Ainsworth & Tenaillon, 2016; Yao *et al*. 2015), in which case approaches based on gene functions may not be powerful.

### QTLs for desired traits in Penicillium roqueforti

We observed a large diversity of phenotypes within each of the generated progenies compared to current commercial strains, and an even greater diversity across the five progenies. We detected pleiotropic regions, acting on multiple metabolism traits, which could facilitate or impair the selection for desired combinations of lipolysis, proteolysis and color. The presence of additional QTLs elsewhere in the genome suggests that a finer selection could be possible for one of the traits without affecting the others. Therefore, our study paves the way for strain improvement and diversification in cheese-making strains.

Toxin production is a concern in *P. roqueforti*, in particular the production of PR toxin and roquefortine C. The low toxin concentrations found in cheese and the long history of safe consumption of blue cheeses, combined with the recent discovery that both Roquefort and non-Roquefort populations are unable to produce PR toxin, indicate the innocuity of commercial strains in cheese production (Crequer *et al*. 2024). Nevertheless, selecting strains with even lower toxin production would be of major interest. We found several QTLs with effects on extrolite production. In particular, QTLs in the biosynthesis gene clusters of PR toxin, roquefortine C and MPA indicated that the corresponding non-Roquefort alleles decreased extrolite production. This suggests selection for lower toxicity in the domesticated non-Roquefort lineage. In addition, lower production of several extrolites was associated with the duplication of the R1 region. Selection of these alleles could thus further improve the safety of *P. roqueforti* use in cheese production or for new products such as vegan cheese. While toxin production is detrimental for cheese safety, other extrolites can be interesting to produce for medical applications: for example, MPA is an immunosuppressant used for preventing transplant organs rejection (Matas *et al*. 2013), and andrastin A, a potential antitumor compound (Ge *et al*. 2009; Matsuda, Awakawa & Abe, 2013). Our study shows large positive transgression for the production of these extrolites and multiple QTL regions. These results are highly promising to increase their production level in the pharmaceutical context.

We detected QTLs for color as well as for protein and lipid degradation. In blue cheeses, lipolysis and proteolysis by *P. roqueforti* influence texture, taste and the production of aromatic compounds, the latter being mainly influenced by lipolysis (Cantor *et al*. 2004). Currently, the texture and the amount of aromatic compounds in the final product is mostly controlled by the length of the ripening period. A longer ripening time increases the duration of lipolysis and proteolysis in cheeses, leading to a softer texture and the production of more pronounced aromatic compounds in the final product (Cantor *et al*. 2004). Although the general consumer demand is moving towards milder products (Michel Place, L.I.P. S.A.S. pers. com.), producers aim to develop a range of products with varying degrees of flavor to meet the diversity of consumer preferences (*i.e.* product segmentation). For example, the Carles firm producing Roquefort PDO cheeses generated a new strain (called “Maxime*”*) producing milder cheeses, with lighted blue color, called “Elegance” (https://www.professionfromager.com/salon-virtuel/france/occitanie/article/carles). Another interest of producers is the visual aspect of the cheese, which depends on the initial color of the spores and the ability of the strain to maintain this color in conditioned atmospheres (Fairclough, Cliffe & Knapper, 2011). We identified QTLs for these traits, indicating that selection for strain improvement and diversification should be possible, for example the selection of strains with different colors, and proteolysis or lipolysis activities.

Breeding could thus be performed in *P. roqueforti* using crosses between different populations and then back-crosses to retain only the desired new traits while purging deleterious mutations. It would thus be possible to use the QTLs identified here for marker-assisted selection, as done in *Saccharomyces cerevisiae* for malic consumption (Vion *et al*. 2021). As *Penicillium roqueforti* is haploid, there is no issue of inbreeding depression and selection is particularly efficient, as there is no dominance effect of masking alleles. Marker-assisted selection is all the more useful in *P. roqueforti* as the assessment of cheese features requires a long and time-consuming process of cheese making with multiple replicates in controlled environments (Caron *et al*. 2021).

## Conclusion

In this study, we managed to generate F1 offspring from different crosses, including one between parents from the two domesticated cheese populations, and obtained high variability in phenotypes in progenies. This is crucial given the very low diversity currently available in each of the Roquefort and non-Roquefort cheese populations. Despite genomic rearrangements and segregation biases due to deleterious alleles, the QTL approach identified regions involved in important traits for cheese making. The high heritabilities and the QTLs identified for all traits but growth indicate that selection should be efficient for strain improvement. Segregation biases, linkage drag, pleiotropy and genomic rearrangements may however reduce the efficacy of selection.

Further efforts could increase the number of offspring and use the Termignon cheese population recently discovered in non-inoculated French cheeses produced at a small scale in the Alps, as this population is genetically and phenotypically differentiated from the two other cheese populations (Crequer *et al*. 2023). With a better understanding of the demands of consumers and cheese producers, association genetics could become a very promising tool for the improvement of *P. roqueforti* strains, increasing blue cheese quality and diversity. Few studies have addressed these questions in the domestication of fungi so far, beyond yeasts and the button mushroom (Foulongne-Oriol *et al*. 2012; Imbernon *et al*. 1996; Nguyen *et al*. 2022; Kessi-Pérez, Molinet & Martínez, 2020; Eder *et al*. 2018; Wilkening *et al*. 2014; Swinnen, Thevelein & Nevoigt 2012; Liti & Louis, 2012; Steinmetz *et al*. 2002). From a fundamental point of view, identifying the genomic regions controlling traits of interest to humans helps understand the genetic mechanisms behind the adaptation of organisms to their environment. In particular, our findings of similar QTLs between independently domesticated *P. roqueforti* lineages, with pleiotropic effects, indicate convergent adaptation targeting major regulators.

## Materials and methods

### Progeny production

The production of progenies was obtained following the protocol in Ropars *et al*. (2014) with some modifications. Six *P. roqueforti* strains (Table 1) were selected for their fertility as observed in a previous study (Ropars *et al*. 2016) and based on their phenotypic differences estimated from proteolysis and lipolysis measurements (Dumas *et al*. 2020). They were crossed in all possible pairs given their mating types, either MAT1-1 (three strains) or MAT1-2 (three strains) (Table 1).

For each cross, three Petri dishes were inoculated on an oat medium supplemented with biotin after sterilization (6.4 μg/L; Böhm *et al*. 2012). On each Petri dish, 5 μL of an uncalibrated spore suspension was used to inoculate agar sections at opposite sites on the Petri dish, located perpendicular to the sections inoculated with aliquots of conidia of the opposite sexual type (Figure 5 A; O’Gorman, Fuller & Dyer, 2009). Petri dishes were sealed with food-grade plastic film and incubated upside down at 19°C in the dark. The contact zone between strains (Figure 5 A) was examined with a binocular loupe and an optical microscope regularly for four to six weeks to check for the presence of cleistothecia (sexual structures). When cleistothecia were found, three to five of them were sampled, crushed with tweezers, and observed under an optical microscope with lactophenol cotton blue dye solution for ascospore detection (Figure 5 B and C). We took the pictures with a Nikon DS-Fi2 camera, on an inverted optical microscope with apodized phase contrast (Nikon France, Champigny-sur-Marne, France). When ascospores were identified, all the cleistothecia of the corresponding Petri dish were collected in a 0.05% Tween-20 water suspension (P1379-100mL Sigma Aldrich, Saint Quentin Fallavier, France). Because cleistothecia are covered with conidia derived from asexual reproduction, we developed a treatment to kill them before releasing ascospores from cleistothecia. Approximately 500 μL of the cleistothecia suspension was pooled with 500 μL of a commercial 2.6% sodium hypochlorite solution and gently agitated for 20 seconds; then 1 mL of water was added to stop the sodium hypochlorite action. Cleistothecia were washed twice with one-minute centrifugation at 7,000 rpm, removal of the supernatant and resuspension in Tween 20. Then, we crushed cleistothecia with a sterile piston (Piston Pellet Eppendorf, Montesson, France) and filtered residues with a 40 μm filter (cell screen, Greiner, Les Ulis, France). Single-ascospore isolation was carried out with a dilution method on malt extract agar Petri dishes (MEA, Galloway and Burgess 1952).

**Figure 5:**
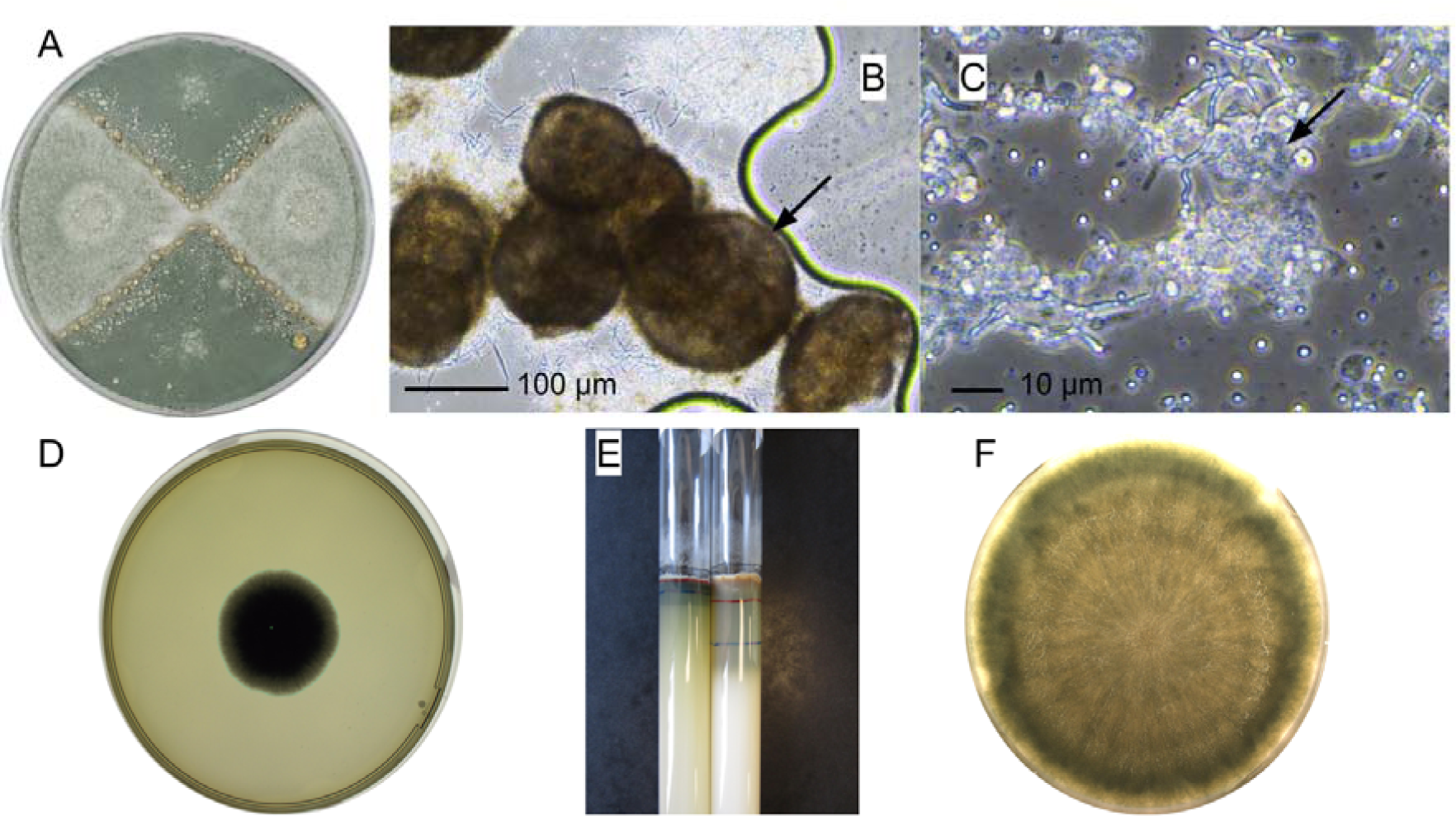
Observations of *Penicillium roqueforti* crosses and examples of phenotypic trait determination. (A) Petri dish with a cross between the LCP06136 (MAT1-1; Roquefort; at the top and bottom) and LCP06173 (MAT1-2; non-Roquefort; at left and right) strains. Sexual structures are formed at the contact zones between the two strains. (B) Cleistothecia (sexual structures of *Penicillium roqueforti*) shown by black arrow. (C) Asci containing ascospores shown by the black arrow. (D) Petri dish used for estimating colony growth rate, by counting the number of pixels occupied by the fungal colony (one offspring grown on malt medium for 120 hours). (E) Two lipolysis test tubes showing the lysis dynamics, for two different strains, with marks on the tubes indicating the limit of lipid medium degradation at different times (blue, red and blue marks, drawn at 0, 7 and 14 days, respectively). (F) Petri dish on which an offspring grew on raw sheep milk medium during 13 days for color measure.

### Recombinant offspring detection

We chose five fertile crosses for further analyses (Table 1). Genomic DNA was extracted from fresh mycelium and conidia suspension after single-ascospore isolation and growth for five days on MEA media using the Nucleospin soil kit (Macherey-Nagel, Düren, Germany). For checking that isolated spores were recombinant offspring and not asexual conidia, we used 11 microsatellite markers (Ropars *et al*. 2014) labeled with fluorescence, out of which between five and nine were polymorphic depending on the cross. These markers were amplified with the Multiplex PCR kit (Qiagen, Les Ulis, France) using a touchdown program with an initial denaturation of 15 min at 95°C, 35 cycles of 30 s at 94°C, a decrease of 1°C every 90 s from 60 to 50°C, and 60 s at 72°C. The PCR program ended with a final 30 min extension step at 60°C. Genotyping by capillary fractionation electrophoresis was performed at INRAe Clermont-Ferrand (INRAe Platform GENTYANE, Clermont-Ferrand, FRANCE). The profiles obtained were analyzed with the GENEMAPPER v4.0 software (Applied Biosystem, Villebon-sur-Yvette, France) to detect recombination between parental genotypes. Only strains with recombinant genotypes were retained, *i.e.* carrying alleles from one parent at some markers and alleles for the other parent at other markers.

### Genome sequencing, assembly and analysis

We generated long-read-based genome assemblies by sequencing the genomes of the LCP06136 parental strains, as well as 24 F1 offspring, with Oxford Nanopore MinION technology with an R9 flow cell, a high-quality genome assembly of the other parental strain (LCP06133) being available (Crequer *et al*. 2023). We also generated genome assemblies for the other parents, based on Illumina sequencing and assemblies guided by the Oxford Nanopore assemblies of the LCP06136 and LCP06133 parents.

DNA was extracted from mycelium and conidia with the Nucleospin soil kit for the progeny and with the NucleoBond High Molecular Weight DNA kit (Macherey-Nagel, Düren, Germany) for the parental strain LCP06136, with mechanical disruption of about 30 mg of lyophilized mycelium with two tungsten beads (3 mm diameter) for 5 min at 30 Hz. The Nanopore library was prepared with the SQK-LSK109 ligation sequencing kit, starting with 1.5 µg DNA, and sequencing was performed in-house with a MinION combined with a MinIT (version 19.06.9), a 72 hour run, and with the Fastcalling basecalling algorithm. The genome of the LCP06136 strain was sequenced alone in a R9 flow cell, whereas the 24 offspring genomes were sequenced in a 12-plex with the Rapid Barcoding Sequencing kit SQK-RBK004. We assessed run quality using Porechop v0.2.3_sequan2.1.1 (Wick *et al*. 2017), and when necessary, demultiplexed the Nanopore raw reads with the default parameters.

*De novo* assemblies of the genomes were constructed from both Illumina and Nanopore reads. For the LCP06136 parental genome, the raw Nanopore reads were trimmed and assembled with Canu v1.8 (Koren *et al*. 2017) with the option genomeSize=28m. The assembly obtained with Canu was polished twice with Illumina reads (Dumas *et al*. 2020) using Pilon v1.24 (Walker *et al*. 2014) with the default settings. For each round of polishing, Bowtie2 <u>(Langmead and Salzberg, 2012;</u> Langmead *et al*. 2019) was used to align the Illumina trimmed reads with the assembly for polishing with a maximum length (-X) of 1000bp. The Illumina reads were first trimmed with Trimmomatic v0.36 (Bolger, Lohse & Usadel, 2014) with the following options: ILLUMINACLIP:TruSeq3-PE.fa:2:30:10 LEADING:10 TRAILING:10 SLIDINGWINDOW:5:10 MINLEN:50. Redundant contigs were removed based on a self-alignment using NUCmer version 3.1 (Kurtz *et al*. 2004). For other parental strains (LCP06037, LCP06039, LCP06043, LCP06059), available Illumina reads (Dumas *et al*. 2020) were trimmed with Trimmomatic v0.36 (Bolger et al 2014) the same way as previously described, and assembled with SOAPdenovo2 v2.04 (Luo *et al*. 2012) using the corresponding default size 23 k-mer length. The assemblies obtained were then scaffolded using Ragout v2.3 with the LCP06136 and the LCP06133 genomes as references (Kolmogorov *et al*. 2018). For the 24 offspring genomes, the raw Nanopore reads were trimmed and assembled both with Canu v1.8 (Koren *et al*. 2017) and with Flye version 2.9 (Kolmogorov *et al*. 2019), and then merged using Quickmerge (Solares *et al*. 2018). The merged assemblies obtained were polished once with a concatenation of Illumina reads of the parental strain LCP06136 and LCP06133 (Dumas *et al*. 2020) in the same way as for the LCP06136 parental genome. The obtained assemblies were scaffolded using Ragtag v2.1.0 (Alonge *et al*. 2022). We evaluated the quality of the final assemblies with Quast v5.0.2 (Gurevich *et al*. 2013) and their completeness with BUSCO v5.3.2 (Manni *et al*. 2021) using the eurotiales_odb10 lineage dataset. Based on the quality and completeness, we selected 11 out of the 24 offspring strain assemblies for further analyses.

Transposable elements (TEs) were annotated using the REPET package (https://urgi.versailles.inra.fr/Tools/REPET). The TEdenovo pipeline (Flutre *et al*. 2011) was used to detect repeated elements in LCP06136 and LCP06133 (https://doi.org/10.57745/SIP7CH) genomes, and to provide consensus sequences. These consensus sequences were classified with PASTEC v1.3 (Hoede *et al*. 2014), based on the Wicker hierarchical TE classification system (Wicker *et al*. 2007) and manually curated. The resulting bulk library of consensus sequences was then used to annotate TE copies in the six whole genomes (namely, LCP06136 and LCP06133, LCP06037, LCP06039, LCP06043, LCP06059) using the TEannot pipeline (Quesneville *et al*. 2005). Repeat-induced point mutations (RIP) in assemblies were detected using the web-based tool The RIPper (Van Wyk *et al*. 2019).

All final assemblies were formatted (contigs being renamed and ordered in descending order of size), and repeats of the LCP06133 and LCP06136 parental genomes were masked with RepeatMasker v4.1.2 (Smit et al 2013) after *de novo* repeat detection using RepeatModeler v2.0.2 (Smit and Hubley 2008). We performed gene annotation on the masked assemblies of the parental strain LCP06136 using the Funnannotate v1.8.9 pipeline (Palmer and Stajich 2020), with Braker v2.1.6 (Brůna *et al*. 2021; Hoff et al. 2016; Hoff *et al*. 2019; Stanke *et al*. 2006; Stanker *et al*. 2008), which uses a combination of the *ab initio* gene predictors Augustus and GeneMark-ET (Lomsadze et al. 2014) with NCBI blast (Altschul *et al*. 1990) and blast+ (Camacho *et al*. 2009). We ran BRAKER twice, first using the BUSCO dataset of proteins eurotiales_odb10 (Manni *et al*. 2021) with the ProtHint pipeline (Brůna et al. 2020; Buchfink et al. 2015; Gotoh et al. 2014; Iwata and Gotoh, 2012; Lomsadze *et al*. 2005), and then using the RNA-seq dataset from Punt *et al*.(2020) and mapped using STAR v2.5.4b (Dobin *et al*. 2013) with the default settings. We combined the results of the two BRAKER runs with TSEBRA (the Transcript Selector for BRAKER; https://github.com/Gaius-Augustus/TSEBRA).

We studied synteny between assemblies by investigating collinearity of one-to-one orthologs along whole genome alignments, using nucmer v3.1 (Kurtz *et al*. 2004), with 500pb as the minimum match length. We visualized synteny between genomes by plotting one-to-one ortholog links using Circos v0.69-6 (Krzywinski *et al*. 2009).

### Phenotyping

We selected four traits for their applied interest in cheesemaking that are easily measurable with precision and at high throughput, for which we assessed the phenotypes in all the isolated offspring obtained for the five analyzed crosses. The growth rate was measured using a ScanStation (Intersciences, Mourjou, France), an incubator for temperature-controlled growth (set at at 22.5°C) taking pictures regularly for each of 100 Petri dishes as well as analyzing images. We spread 5 μL of standardized spore suspension (250 spores/inoculation) on each Petri dish, containing malt medium (Biokar BK045 HA) at 15 g.L^-1^ in source water (Cristalline, Chelles, France), sterilized at 120°C during 15 minutes. The ScanStation took pictures of the Petri dishes right side up on a black background, for five days every 30 minutes, *i.e.* 240 measures per strain (Figure 5 D). The ScanStation estimated the colony area and diameter for each picture, by counting the number of pixels corresponding to the fungal colony. Because the colony diametral growth was near linear when plotted against days (Supplementary Figure 6), we used, as a measure of growth rate, the slope of the least square linear regression computed with the *lm* function of the base package R software v4.3.2 (http://www.r-project.org/). The coefficients of determination (r^2^), measuring the fit of the data to a line, are presented in the Supplementary Table 5 and the linear regression of an offspring with a median r^2^ in the Supplementary Figure 6. To ensure the robustness of the linear fit, strains with coefficients of determination below 0.98 were not considered in the QTL analysis of this trait (21 of 885, *i.e.* 2.4% of strains filtered out).

The colony color was measured after growth on raw sheep milk powder medium (Biocoop, Paris, France) at 21% in source water, sterilized at 105°C for five minutes, and mixed with 1.7% agar in source water, sterilized at 121°C for 15 minutes. After 13 days of growth, two pictures were taken by the ScanStation under standardized conditions of light and on a white background (Figure 5 F). We recorded for each strain the colony color decomposition using RGB (red, green and blue) and HSB (hue, saturation, and brightness) with ImageJ v1.52n (Schneider 2012; Supplementary Table 6). In the RGB system, the levels of red, green and blue are represented by a range of integers from 0 to 255 (256 levels for each color). In the HSB system, hue ranges from 0 to 360 (0 is red, 120 is green, 240 is blue), saturation and brightness from 0 to 100% (0% saturated is neutral grey, 100% saturated is the full color ; 0% brightness is black, 50% brightness is normal, 100% brightness is white). In order to obtain uncorrelated components from the RGB data, we divided each value by the sum of the three components. For each of these parameters, we recorded average values across all the colony pixels for each strain. For QTL detection, we chose *a posteriori,* among these color parameters, those whose distributions and heritabilities were optimal for QTL analysis (*i.e.* largest and uncorrelated variance); we thus retained the RGB proportion, as well as HSB.

The lipolytic and proteolytic activities of *P. roqueforti* strains were measured *in vitro* following Dumas *et al*. (2020): for each strain, 50 μl of standardized spore suspensions (ca. 2,500 spores/inoculation) were inoculated at the top of a test tube containing agar and tributyrin for lipolytic activity measure (10 mL.L^-1^, ACROS Organics) or semi-skimmed cow milk for the proteolytic activity measure (40 g.L^-1^, Casino). The lipolytic and proteolytic activities were estimated by the degree of compound degradation, which changes the media from opaque to translucent. We measured the distance between the initial mark and the hydrolyzed, translucent front, after 7, 14, 21 and 28 days of growth at 20°C in the dark (Figure 5 E). As lipolysis curves were nearly linear when plotted against days, we estimated the lipolysis rate with a linear regression computed with the R base package function *lm*. Proteolysis profiles appeared less linear but with no clear sigmoid nor other regression patterns, so we also chose to fit linear models for more simplicity. The coefficients of determination (r^2^), measuring the fit of the data to a line, are presented in the Supplementary Table 5 and an example of a linear regression of an offspring with a median r^2^ in the Supplementary Figure 6.

For extrolite production, we grew fungal cultures in 24-well sterile microplates containing 2 mL of yeast extract sucrose (YES) agar medium buffered at pH 4.5 with phosphate-citrate buffer and characterized by a high C/N ratio to favor extrolite production as previously described (Frisvad & Filtenborg, 1983). For each strain, 1 µL of a calibrated spore suspension (10^6^ spores.mL^-1^) prepared from a 7-day culture was inoculated in the centre of the well. Two replicates per strain were performed for extrolite analyses. The plates were incubated at 25°C in the dark for seven days and then stored at - 20°C until extrolite analysis. For extrolite extractions, we used an optimized “high-throughput” extraction method (Lo *et al*. 2023; Crequer *et al*. 2024). Briefly, 2g-aliquots (the entire YES culture obtained from a well) were homogenized after thawing samples with a sterile flat spatula then 12.5 mL of acetonitrile (ACN) supplemented with 0.1% formic acid (v/v) was added, samples were vortexed for 30 sec followed by 15 min sonication. Extracts were again vortexed before being centrifuged for 10 min at 5000g at 4°C. The supernatants were directly collected and filtered through 0.45 µm PTFE membrane filters (GE Healthcare Life Sciences, UK) into amber vials and stored at -20°C until analyses. Extrolite detection and quantification were performed using an Agilent 6530 Accurate-Mass Quadropole Time-of-Flight (Q-TOF) LC/MS system equipped with a Binary pump 1260 and degasser, well plate autosampler set to 10°C and a thermostated column compartment. Filtered 2 µL aliquots were injected into a ZORBAX Extend C-18 column (2.1×50 mm and 1.8 µm, 600 bar) maintained at 35°C with a flow rate set to 0.3 mL.min^-1^. The mobile phase A contained milli-Q water + 0.1% formic acid (v/v) and 0.1% ammonium formate (v/v) while mobile phase B was ACN + 0.1% formic acid. Mobile phase B was maintained at 10% for 4 min, followed by a gradient from 10 to 100% for 18 min. Then, mobile phase B was maintained at 100% for 5 min before a 5-min post-time. Samples were ionized in positive (ESI+) electrospray ionization modes in the mass spectrometer with the following parameters: capillary voltage 4 kV, source temperature 325°C, nebulizer pressure 50 psig, drying gas 12 L.min^-1^, ion range 100-1000 m/z. Target extrolite characteristics used for quantifications are given in Supplementary Table 7 and included commercially available extrolites produced by *Penicillium* species, namely andrastin A, eremofortins A & B, (iso)-fumigaclavin A, mycophenolic acid and roquefortine C. Andrastin A, eremofortins A & B and (iso)-fumigaclavin A standards were obtained from Bioviotica (Goettingen, Germany), while all others were from Sigma-Aldrich (St Louis, MO, USA). All stock solutions were prepared in dimethyl sulfoxide (DMSO) at 1 mg.mL^-1^ in amber vials. As the PR toxin was not commercially available, a previously produced purified stock solution (Gillot *et al*. 2017) without known concentration was used to ensure MPA (mycophenolic acid) and PR toxin were separated (as they have the same mass) as well as linearity of PR toxin quantification. For these analyses, metabolite identification was performed using both the mean retention time ± 1 min and the corresponding ions listed in Supplementary Table 7. We used a matrix-matched calibration curve (r^2^ >0.99 for all extrolites except 2 >0.96) to confirm linearity the relation between signal area and extrolite concentration with final concentrations ranging from 1 to 10000 ng.mL^-1^ according to the target metabolite and method performance was carried out as previously described (Gillot *et al*. 2017).

### Statistical analysis on phenotypes

For phenotype distributions, we performed tests of normality with the *shapiro.test* function of the base package of R software v4.3.2 (http://www.r-project.org/). We ran tests for unimodality with the *dip.test* function of the Diptest package v0.75-7 (Maechler 2016) and tests for bimodality with the *bimodality_coefficient* function of the Mousetrap package v3.2.1 (Kieslich *et al*. 2019) with a threshold of 0.555 according to Pfister *et al*. 2013. We performed the trimodality and amodality tests with the *is.trimodal* and *is.amodal* functions, respectively, of the LaplacesDemon v16.1.6 package (Hall *et al*. 2021). We computed the average heterosis as ((F1 - MP) / MP) * 100, F1 and MP being the trait mean values of the progeny and the parents, respectively.

We computed the matrix of Pearson and Spearman correlation coefficients with the R stats package v4.3.2 (http://www.r-project.org/). We produced the violin plots using vioplot v0.4.0 (Adler *et al*. 2022), ggplot2 v3.4.3 (Wickham 2016) and reshape2 v1.4.4 (Wickham 2007) R packages. We performed the principal component analysis (PCA) using the function *PCA* from the R base package, assigning trait means to missing values. Plots were drawn using FactoMineR v2.8 (Lê *et al*. 2002), factoextra v1.0.7 (Kassambara and Mundt 2020) and corrplot v0.92 (Wei and Simko 2021) packages.

### Phenotype heritability

Because we could not perform replicated measurements of phenotypes for each strain for lipolysis, proteolysis, growth rate and color, we only estimated the narrow sense heritability (h^2^) of these phenotypes across the five progenies by estimating the slope of linear regressions of the five F1 progenies mean trait values on the mean values of the parental pairs using the *lm* function from the R base package. Extrolite production was determined for three crosses (SxN, RxN and RxL), with two replicates for each strain; we therefore estimated broad-sense heritability (H^2^) for these phenotypes for each cross based on variability among replicates, in addition to narrow-sense heritability as for the other phenotypes. Broad sense heritability was determined by performing a one-way ANOVA with the *lm* function of the stats v4.2.1 package in R.

### Genotyping based on indels

We genotyped 389 offspring from the cross RxN between LCP06136 (Roquefort; MAT1-1; https://www.ebi.ac.uk/ SAMEA103939766) and LCP06133 (non-Roquefort; MAT1-2; SAMEA103939763) with 200 markers with a polymorphism of amplification size between the two parental strains, due to the presence of indels (Supplementary Table IND). Primers were designed (i) to yield an amplicon size between 150 to 410 bp and a difference ranging from six to 20 bp between alleles, (ii) to be compatible in multiplex and (iii) to be evenly distributed along the assembly of the reference genome available at the beginning of the study (FM164; EMBL accession numbers HG792015 to HG792062, Cheeseman & Ropars *et al*. 2014). Indels were detected after a whole assembly alignment (blast v2.9.0; Altschul *et al*. 1990) of the two parental strains with the gap and extension penalties set to 1. After adding 250 base pairs upstream and downstream from the hit position, we performed a multi-alignment using MAFFT (Katoh 2002) with the whole genomes of the parental strains of the other crosses. We only kept a consensus sequence generated using consambig (Rice et al. 2000) and designed primers using Primer3 (Untergasser *et al*. 2012).

To reduce the cost of fluorescence primers, we only used four universal fluorescent primers (M13(-21), D8S1132f, D12S1090f, DYS437f), labeled with different fluorescent dyes (FAM, ATTO550, ATTO565, YAKIMA Yellow; Eurofins, Ebersberg, Germany). The specific primers attached during PCR to universal fluorescent primers thanks to a tail added to each specific primer matching an edge of the universal primers (Schuelke 2000; Missiaggia and Grattapaglia 2006). For each locus, the PCR amplification thus required three primers: the forward fluorescent universal primer, the forward specific primer with a 5’ tail corresponding to the chosen universal primer and the regular specific reverse primer. The 200 markers were genotyped in 25 multiplex groups of 8 locus organized according to their amplicon size and their color fluorescent dye (Supplementary Table 8). For each multiplex, two separate sets of quadruplex PCR reactions were performed to minimize the interaction between primers and then pooled into octoplexes for the genotyping step. The PCR mix contained 0.16 µM of each of the fluorescent universal primer and of the reverse specified primer and 0.04 µM of the 5’ tail forward primer in a final 15 µl reaction volume (2x QIAGEN Multiplex PCR Master Mix with 3 mM Mg^2+^, 10x primer mix with 1.6 or 0.4 µM of each primer, 3 µl of DNA diluted 50 fold) using the same PCR cycle conditions as for the microsatellites loci previously described in the recombinant offspring detection paragraph, except that it started with a hybridization temperature of 62°C instead of 60°C. Genotyping and analysis were also done as previously described in the recombinant offspring detection paragraph.

### Genotyping based on GBS

For all five crosses, we genotyped the obtained progeny (1073 offspring in total) at the AGAP CIRAD platform with genotyping-by-sequencing using the *Ape*KI enzyme. DNAs were purified after digestion on QIAquick columns (Qiagen, Les Ulis, France) and their qualities were controlled using a TapeStation instrument (Agilent Technologies, Les Ulis, France). Sequencing was performed on an Illumina Hiseq 4000 (2×150 bp). The following numbers of recombinant offspring were genotyped by GBS: 384 for the cross RxN, 157 for the cross RxS, 185 for the cross RxL, 176 for the cross SxN and 171 for the cross LxN.

Demultiplexed reads were mapped to the LCP06136 high-quality genome for the crosses RxL and RxS and to the LCP06133 high-quality genome for the crosses RxN, LxN and SxN using *Bowtie2* version 2.3.4.1 (Langmead and Salzberg 2012; Langmead *et al*. 2019). In *Bowtie2,* we set the maximum length (-X) to 1000 and used the preset “very-sensitive-local”. We used SAMtools v1.7 (Danecek *et al*. 2021) to sort and filter out duplicate reads and reads with a mapping quality score above ten for SNP calling. Single nucleotide polymorphisms (SNPs) were called using the GATK v4.1.2.0 Haplotype Caller <u>(McKenna *et al*. 2010</u>), generating a gVCF file per strain with option ploidy 2 to detect potential duplicated regions in progenies, as we had found indels with two alleles in some offspring. GVCFs were combined using GATK CombineGVCFs, genotypes with GATK GenotypeGVCFs, and biallelic SNPs were selected after filtration using GATK SelectVariants. We filtered SNPs using GATK VariantFiltration and options QUAL <30, DP < 5, QD < 20.0, FS > 60.0, MQ < 40, 0.05<AF<0.95, SOR > 3.0, MQRankSum < -12.5, ReadPosRankSum < -8.0, DP>5.0 and GQ<10. We generated the genotype matrix based on the VCF file with *Bcftools* v1.11 (Li 2011; Danecek *et al*. 2021). Markers with no data or with no differences between the two parental genomes were filtered out. To reconstruct offspring genotypes, we performed SNP calling in the diploid mode in order to detect double copies of translocated regions. Markers corresponding to the translocated regions were in fact called as heterozygous in some offspring; they were therefore placed in offspring at the two parental locations, which were each assigned the corresponding parental allele.

After performing crosses and isolating ascospores, we detected sectors of two slightly different morphotypes in subsequent cultures of the parental strain LCP06039 aiming at phenotype measurements. However, genetic data indicated that the cross was performed with a pure strain and not a mixture: (i) the indel genotypes of the two morphotypes were assessed and were identical, and (ii) the GBS genotypes of the two morphotypes were not typed directly, but we did not detect genome-wide segregation distortion in the progeny in GBS data for this cross, while a cross involving a mixture of strains with different genotypes should result in over-representation of the alleles by the pure parent; on the contrary, the segregation distortions detected in some genomic regions corresponded to over-representations of the strain showing two morphotypes rather than its under-representation. We therefore considered that a single strain contributed to the cross and that the two morphotypes were the result of plasticity or a few somatic mutations in a single strain. We therefore built the genetic map and ran a QTL analysis for the LxN cross. For analyses involving parental traits, and therefore potentially affected by the presence of two morphotypes (*i.e.* for heritability estimates and transgression detection) we computed estimates using all three possible values for the LCP06039 parent strain phenotype: the means of the trait values for the two morphotypes or the value of one of the two morphotypes. This did not affect conclusions as the two morphotypes were actually very close for all trait measures.

### Experiment on the stability of genotypes across replication

As we identified a genomic region with two alleles present in multiple offspring in the RxN cross with associated QTLs, we wanted to check whether this particular genotype was stable during strain culture and across multiple replication steps. We therefore cultivated, on malt agar, the two parental strains from the RxN cross and the 11 offspring for which genomes were sequenced with Nanopore, six with alleles of both parents for this genomic region, four with the LCP06136 parental allele and one with the LCP06133 allele. We cultivated the strains for one week on malt agar and replated them on a new plate by transplanting some conidia from the edge of the colonies. We performed 19 replication steps. In the end, we re-genotyped the lines with the six indel markers present in the rearranged region.

### Linkage map and QTL detection

Obtained offpring phenotypes are presented in Supplementary Table 9 and offpring genotype are presented in Suplementary Tables 10, 11, 12, 13 and 14 for SxN, LxN, RxN, RxS and RxL crosses respectively. The genetic map and statistical tests for QTL association were performed using the R software v3.5.1 (http://www.r-project.org/) with the ASMap v1.0-5 package (Taylor and Butler 2017) for the genetic map and qtl v1.60 package (Broman *et al*. 2003) for statistical tests. For the genetic map, we used unique segregating markers with less than 10% missing data and markers with segregation distortion with significant Chi Square p-values adjusted with Bonferroni corrections. Individuals with more than 30% of missing data were filtered out (Table 1), concerning up to 14% of the offspring of a cross. Linkage groups and marker ordering were determined with the *mstmap* function of AsMap package, with parameter “bychr” set as FALSE and a p.value of 10^-12^. Markers with less than 20% missing data and less than 95% of one parental allele were then reintegrated in linkage groups with the *pull.cross* function with “max.rf” and “min.lod” parameters set at 0.1 and 20, respectively. We reconstructed marker order in linkage groups with the *mstmap* function, the parameter “bychr” set as TRUE, and a p.value< 10^-12^. We estimated genetic distances between markers with the AsMap *quickEst* function and the Kosambi’s mapping function. We compared the obtained order to the reference genome order using the *compareorder* function of the qtl package with error probability set at 0.005. We changed the order only if it improved the LOD score and minimized the linkage group length. We represented the relationships between chromosome physical positions and genetic maps using ALLMAPS in JCVI utility libraries v1.3.6 (Tang *et al*. 2015), adapted to include genetic features.

We performed single QTL detection using the *scanone* function of the qtl package v1.6 with the threshold at 5% being estimated with 1000 permutations. We used the *addqtl* function of the qtl package to mask the first set of QTLs and thus detect additional QTLs that may have been hidden by the main ones. We estimated confidence intervals for QTL using the *lodint* function of the qtl package with default parameters. We identified QTL interactions using the *scantwo* function with the Haley-Knott method using a threshold at 5% estimated by 1,000 permutations. We performed analyses of variance (ANOVAs) using a linear model with the *fitqtl* function of the qtl package, keeping interactions between loci only when they were significant in both *scantwo* and the full model AVOVAs. We extended QTL confidence intervals including the first or last marker of a linkage group to the start or end of the corresponding chromosome. For comparisons of QTL interval positions between crosses, we transposed the intervals determined with the positions on the Roquefort parent genomes (for RxL and RxS crosses) onto the non-Roquefort genome. To do this, we extracted genome sequences using bedtools getfasta v2.30.0 (Quinlan and Hall 2010) and mapped the extracted sequences onto the non-Roquefort parent genome using minimap2 v2.17-r941 (Li 2018). The representation of the aligned QTL intervals was constructed using the karyoplotR v1.24.0 package (Gel and Serra 2017).

## Supporting information

Supplementary files containing Supplementary figures 1 to 6

Supplementary file containing supplementary tables 1 to 14

## Acknowledgements

This study has been funded by the ANR-19-CE20-0002-02 Fungadapt ANR, ERC Genomefun 309403 Stg, ERC Blue Proof of Concept, Fondation Louis D (French Academy of Sciences) grants, the LIP SAS and the ANRT (Association Nationale Recherche Technologie). TC and ECr acknowledge the E3GP3 network for facilitating collaborations and skills transfer from MF laboratory. We acknowledge the staff at the GENTYANE genotyping platform (INRAe GDEC, Clermont-Ferrand, France) for microsatellite and indel genotyping and the AGAP CIRAD platform for GBS sequencing. We gratefully acknowledge Intersciences for the loan and set-up of the ScanStation (https://www.interscience.com/en/products/real-time-incubator-and-colony-counter/). We thank Jeanne Ropars for her help in obtaining the progeny and Jean-Philippe Vernadet for help with bioinformatic analyses.

## Author contribution

TG, TC, AB, DR, CC and MP designed the study. TG, DR, ECo and MP acquired funding. TC, AS and SLP performed crosses, isolated strains and extracted DNAs, supervised by TG. TC, MLP and SB performed experiments for phenotyping lipolysis, proteolysis and growth rates supervised by CC, DR and MP. TC, ASn and SLP performed genotyping and genome sequencing, supervised by TG. ECr and GC performed experiments for estimating the production of extrolites supervised by MC. TC and ASn performed the experiment on the stability of genotypes across replication and the Oxford Nanopore sequencing, supervised by TG. TC, ECr, ASi and RRdlV performed the genomic analyses, supervised by TG, MFO and AB. TC, TG and ECr wrote the manuscript. All authors revised the manuscript.

## Data availability

All accession numbers will be available upon manuscript acceptance.

## References

Adamczyk, K., Pokorska, J., Makulska, J., Earley, B., & Mazurek, M. (2013). Genetic analysis and evaluation of behavioural traits in cattle. Livestock Science, 154, 1–12. 10.1016/j.livsci.2013.01.016

Adler, D., Kelly, T., Elliott, T., & Adamson, J. (2022). *vioplot: Violin plot.* (0.4.0) [Computer software]. https://github.com/TomKellyGenetics/vioplot

Alexa, A., & Rahnenfuhrer, J. (2016). *topGO: Enrichment Analysis for Gene Ontology* [Computer software]. https://bioconductor.org/packages/release/bioc/html/topGO.html

Alonge, M., Lebeigle, L., Kirsche, M., Jenike, K., Ou, S., Aganezov, S., Wang, X., Lippman, Z. B., Schatz, M. C., & Soyk, S. (2022). Automated assembly scaffolding using RagTag elevates a new tomato system for high-throughput genome editing. Genome Biology, 23(1), 258. 10.1186/s13059-022-02823-7

Altschul, S. F., Gish, W., Miller, W., Myers, E. W., & Lipman, D. J. (1990). Basic Local Alignment Search Tool. Journal of Molecular Biology, 215(3), 403–410. 10.1016/S0022-2836(05)80360-2

Andargie, M., Pasquet, R. S., Gowda, B. S., Muluvi, G. M., & Timko, M. P. (2014). Molecular mapping of QTLs for domestication-related traits in cowpea (*V. unguiculata* (L.) Walp.). Euphytica, 200(3), 401–412. 10.1007/s10681-014-1170-9

Baack, E. J., Sapir, Y., Chapman, M. A., Burke, J. M., & Rieseberg, L. H. (2008). Selection on domestication traits and quantitative trait loci in crop–wild sunflower hybrids. Molecular Ecology, 17(2), 666–677. 10.1111/j.1365-294X.2007.03596.x

Bachlava, E., Tang, S., Pizarro, G., Schuppert, G. F., Brunick, R. K., Draeger, D., Leon, A., Hahn, V., & Knapp, S. J. (2010). Pleiotropy of the branching locus (*B*) masks linked and unlinked quantitative trait loci affecting seed traits in sunflower. Theoretical and Applied Genetics, 120(4), 829–842. 10.1007/s00122-009-1212-1

Bernardes, J. P., Stelkens, R. B., & Greig, D. (2017). Heterosis in hybrids within and between yeast species. Journal of Evolutionary Biology, 30(3), 538–548. 10.1111/jeb.13023

Böhm, J., Hoff, B., O’Gorman, C. M., Wolfers, S., Klix, V., Binger, D., Zadra, I., Pöggeler, S., Dyer, P. S., & Kück, U. (2012). Sexual reproduction and mating-type – mediated strain development in the penicillin-producing fungus *Penicillium chrysogenum*. Proceedings of the National Academy of Sciences, 110(4), 1476–1481. 10.1073/pnas.1217943110/-/DCSupplemental. www.pnas.org/cgi/doi/10.1073/pnas.1217943110

Bolger, A. M., Lohse, M., & Usadel, B. (2014). Trimmomatic: A flexible trimmer for Illumina sequence data. Bioinformatics, 30(15), 2114–2120. 10.1093/bioinformatics/btu170

Bomblies, K., & Doebley, J. F. (2006). Pleiotropic effects of the duplicate maize *FLORICAULA/LEAFY* genes *zfl1* and *zfl2* on traits under selection during maize domestication. Genetics, 172(1), 519–531. 10.1534/genetics.105.048595

Broman, K. W., Wu, H., Sen, Ś., & Churchill, G. A. (2003). R/qtl: QTL mapping in experimental crosses. Bioinformatics, 19(7), 889–890. 10.1093/bioinformatics/btg112

Brůna, T., Lomsadze, A., & Borodovsky, M. (2020). GeneMark-EP+: Eukaryotic gene prediction with self-training in the space of genes and proteins. NAR Genomics and Bioinformatics, 2(2), lqaa026. 10.1093/nargab/lqaa026

Brůna, T., Hoff, K. J., Lomsadze, A., Stanke, M., & Borodovsky, M. (2021). BRAKER2: Automatic eukaryotic genome annotation with GeneMark-EP+ and AUGUSTUS supported by a protein database. NAR Genomics and Bioinformatics, 3(1), lqaa108. 10.1093/nargab/lqaa108

Buchanan, R. L., Golden, M. H., & Whiting, R. C. (1993). Differentiation of the effects of pH and lactic or acetic acid concentration on the kinetics of *Listeria monocytogenes* inactivation. Journal of Food Protection, 56(6), 474–479. 10.4315/0362-028X-56.6.474

Buchfink, B., Xie, C., & Huson, D. H. (2015). Fast and sensitive protein alignment using DIAMOND. Nature Methods, 12(1), 59–60. 10.1038/nmeth.3176

Bucknell, A. H., & McDonald, M. C. (2023). That’s no moon, it’s a *Starship*: Giant transposons driving fungal horizontal gene transfer. Molecular Microbiology, mmi.15118. 10.1111/mmi.15118

Camacho, C., Coulouris, G., Avagyan, V., Ma, N., Papadopoulos, J., Bealer, K., & Madden, T. L. (2009). BLAST+: Architecture and applications. BMC Bioinformatics, 10(1), 421. 10.1186/1471-2105-10-421

Cantor, M. D., Van Den Tempel, T., Hansen, T. K., & Ardö, Y. (2004). Blue cheese. *Cheese: Chemistry*, Physics and Microbiology, 2, 175–198. 10.1016/S1874-558X(04)80044-7

Caron, T., Piver, M. L., Péron, A.-C., Lieben, P., Lavigne, R., Brunel, S., Roueyre, D., Place, M., Bonnarme, P., Giraud, T., Branca, A., Landaud, S., & Chassard, C. (2021). Strong effect of *Penicillium roqueforti* populations on volatile and metabolic compounds responsible for aromas, flavor and texture in blue cheeses. International Journal of Food Microbiology, 354, 109174. 10.1016/j.ijfoodmicro.2021.109174

Cerning, J., Gripon, J.-C., Lamberet, G., & Lenoir, J. (1987). Les activités biochimiques des *Penicillium* utilisés en fromagerie. Le Lait, 67(1), 3–39. 10.1051/lait:198711

Chávez, R., Vaca, I., & García-Estrada, C. (2023). Secondary metabolites produced by the blue-cheese ripening mold *Penicillium roqueforti*; biosynthesis and regulation mechanisms. Journal of Fungi, 9(4), 459. 10.3390/jof9040459

Cheeseman, K., Ropars, J., Renault, P., Dupont, J., Gouzy, J., Branca, A., Abraham, A.-L., Ceppi, M., Conseiller, E., Bensimon, A., Giraud, T., & Brygoo, Y. (2014). Multiple recent horizontal transfers of a large genomic region in cheese making fungi. Nature Communications, 5, 2876. 10.1038/ncomms3876

Cleere, M. M., Novodvorska, M., Geib, E., Whittaker, J., Dalton, H., Salih, N., Hewitt, S., Kokolski, M., Brock, M., & Dyer, P. S. (2024). New colors for old in the blue-cheese fungus Penicillium roqueforti. Npj Science of Food, 8(1), 3. 10.1038/s41538-023-00244-9

Collins, Y. F., McSweeney, P. L. H., & Wilkinson, M. G. (2003). Lipolysis and free fatty acid catabolism in cheese: A review of current knowledge. International Dairy Journal, 13(11), 841–866. 10.1016/S0958-6946(03)00109-2

Coluccio, A., Bogengruber, E., Conrad, M. N., Dresser, M. E., Briza, P., & Neiman, A. M. (2004). Morphogenetic pathway of spore wall assembly in *Saccharomyces cerevisiae*. Eukaryotic Cell, 3(6), 1464–1475. 10.1128/EC.3.6.1464-1475.2004

Conrado, R., Gomes, T. C., Roque, G. S. C., & De Souza, A. O. (2022). Overview of bioactive fungal secondary metabolites: Cytotoxic and antimicrobial compounds. Antibiotics, 11(11), 1604. 10.3390/antibiotics11111604

Cosciani-Cunico, E., Dalzini, E., Ducoli, S., Sfameni, C., Bertasi, B., Losio, M.-N., Daminelli, P., & Varisco, G. (2015). Behaviour of *Listeria monocytogenes* and *Escherichia coli* O157:H7 during the cheese making of traditional raw-milk cheeses from Italian Alps. Italian Journal of Food Safety, 4(2). 10.4081/ijfs.2015.4585

Coton, E., Coton, M., Hymery, N., Jany, J. L., & Mounier, J. (2020). *Penicillium roqueforti*: An overview of its genetics, physiology, metabolism and biotechnological applications. Fungal Biology Reviews, 34(2), 59–73. 10.1016/j.fbr.2020.03.001

Crequer, E., Ropars, J., Jany, L., Caron, T., Coton, M., Snirc, A., Vernadet, P., Branca, A., Giraud, T., & Coton, E. (2023). A new cheese population in Penicillium roqueforti and adaptation of the five populations to their ecological niche. Evolutionary Applications, 1–20. 10.1111/eva.13578

Crequer, E., Coton, E., Cueff, G., Cristiansen, J. V., Frisvad, J. C., De La Vega, R. R., Giraud, T., Jany, J.-L., & Coton, M. (2024). Different metabolite profiles across Penicillium roqueforti populations associated with ecological niche specialisation and domestication [Preprint]. Microbiology. 10.1101/2024.01.12.575369

Cryer, N. C., Butler, D. R., & Wilkinson, M. J. (2005). High throughput, high resolution selection of polymorphic microsatellite loci for multiplex analysis. Plant Methods, 1, 1–5. 10.1186/1746-4811-1-3

Danecek, P., Bonfield, J. K., Liddle, J., Marshall, J., Ohan, V., Pollard, M. O., Whitwham, A., Keane, T., McCarthy, S. A., Davies, R. M., & Li, H. (2021). Twelve years of SAMtools and BCFtools. GigaScience, 10(2), giab008. 10.1093/gigascience/giab008

Dobin, A., Davis, C. A., Schlesinger, F., Drenkow, J., Zaleski, C., Jha, S., Batut, P., Chaisson, M., & Gingeras, T. R. (2013). STAR: Ultrafast universal RNA-seq aligner. Bioinformatics, 29(1), 15–21. 10.1093/bioinformatics/bts635

Doebley, J., Stec, A., & Hubbard, L. (1997). The evolution of apical dominance in maize. Nature, 386(6624), 485–488. 10.1038/386485a0

Dumas, É., Feurtey, A., Rodríguez de la Vega, R. C., Le Prieur, S., Snirc, A., Coton, M., Thierry, A., Coton, E., Le Piver, M., Roueyre, D., Ropars, J., Branca, A., & Giraud, T. (2020). Independent domestication events in the blue-cheese fungus *Penicillium roqueforti*. Molecular Ecology, 33913(January), 451773. 10.1111/mec.15359

Eder, M., Sanchez, I., Brice, C., Camarasa, C., Legras, J.-L., & Dequin, S. (2018). Qtl mapping of volatile compound production in *Saccharomyces cerevisiae* during alcoholic fermentation. BMC Genomics, 19, 166. 10.1186/s12864-018-4562-8

Enyenihi, A. H., & Saunders, W. S. (2003). Large-Scale Functional Genomic Analysis of Sporulation and Meiosis in *Saccharomyces cerevisiae*. Genetics, 163(1), 47–54. 10.1093/genetics/163.1.47

Fairclough, A. C., Cliffe, D. E., & Knapper, S. (2011). Factors affecting *Penicillium roquefortii* (*Penicillium glaucum*) in internally mould ripened cheeses: Implications for pre-packed blue cheeses. International Journal of Food Science & Technology, 46, 1586–1590. 10.1111/j.1365-2621.2011.02658.x

Falconer, D. S., & Mackay, T. F. C. (1995). Introduction to Quantitative Genetics. Pearson Education Limited.

Flutre, T., Duprat, E., Feuillet, C., & Quesneville, H. (2011). Considering Transposable Element Diversification in *De Novo* Annotation Approaches. PLoS ONE, 6(1), e16526. 10.1371/journal.pone.0016526

Fontaine, K., Hymery, N., Lacroix, M. Z., Puel, S., Puel, O., Rigalma, K., Gaydou, V., Coton, E., & Mounier, J. (2015). Influence of intraspecific variability and abiotic factors on mycotoxin production in *Penicillium roqueforti*. International Journal of Food Microbiology, 215, 187–193. 10.1016/j.ijfoodmicro.2015.07.021

Foulongne-Oriol, M., Rodier, A., Rousseau, T., & Savoie, J.-M. (2012). Quantitative trait locus mapping of yield-related components and oligogenic control of the cap color of the button mushroom, *Agaricus bisporus*. Applied and Environmental Microbiology, 78(7), 2422–2434. 10.1128/AEM.07516-11

Fox, J., & Weisberg, S. (2019). An {R} Companion to applied regression (T. Oaks, Ed.). Sage. https://socialsciences.mcmaster.ca/jfox/Books/Companion/

Frisvad, J. C., & Filtenborg, O. (1983). Classification of terverticillate *Penicillia* based on profiles of mycotoxins and other secondary metabolites. Applied and Environmental Microbiology, 46(6), 1301– 1310. 10.1128/aem.46.6.1301-1310.1983

Galagan, J. E., & Selker, E. U. (2004). RIP: The evolutionary cost of genome defense. Trends in Genetics, 20(9), 417–423. 10.1016/j.tig.2004.07.007

Galloway, L. D., & Burgess, R. (1952). Galloway L.D. and Burgess R.(1952) Applied Mycology and Bacteriology 3rd Edition Leonard Hill, London. Pp54 and 57. Applied Mycology and Bacteriology, 3, 54–57.

Ge, H. M., Yu, Z. G., Zhang, J., Wu, J. H., & Tan, R. X. (2009). Bioactive alkaloids from endophytic *Aspergillus fumigatus*. Journal of Natural Products, 72(4), 753–755. 10.1021/np800700e

Gel, B., & Serra, E. (2017). karyoploteR: An R/Bioconductor package to plot customizable genomes displaying arbitrary data. Bioinformatics, 33(19), 3088–3090. 10.1093/bioinformatics/btx346

Gil-Durán, C., Rojas-Aedo, J. F., Medina, E., Vaca, I., García-Rico, R. O., Villagrán, S., Levicán, G., & Chávez, R. (2015). The *pcz1* gene, which encodes a Zn(II)2Cys6 Protein, is involved in the control of growth, conidiation, and conidial germination in the filamentous fungus *Penicillium roqueforti*. PLOS ONE, 10(3), e0120740. 10.1371/journal.pone.0120740

Gillot, G., Jany, J.-L., Coton, M., Le Floch, G., Debaets, S., Ropars, J., López-Villavicencio, M., Dupont, J., Branca, A., Giraud, T., & Coton, E. (2015). (Fifty shades of blueL:) Insights into Penicillium roqueforti morphological and genetic diversity. PLoS ONE, 10(6), e0129849. 10.1371/journal.pone.0129849

Gillot, G., Jany, J.-L., Poirier, E., Maillard, M., Debaets, S., Thierry, A., Coton, E., & Coton, M. (2017). Functional diversity within the *Penicillium roqueforti* species. International Journal of Food Microbiology, 241, 141–150. 10.1016/j.ijfoodmicro.2016.10.001

Gladieux, P., Ropars, J., Badouin, H., Branca, A., Aguileta, G., De Vienne, D. M., Rodríguez de la Vega, R. C., Branco, S., & Giraud, T. (2014). Fungal evolutionary genomics provides insight into the mechanisms of adaptive divergence in eukaryotes. Molecular Ecology, 23(4), 753–773. 10.1111/mec.12631

Gluck-Thaler, E., Ralston, T., Konkel, Z., Ocampos, C. G., Ganeshan, V. D., Dorrance, A. E., Niblack, T. L., Wood, C. W., Slot, J. C., Lopez-Nicora, H. D., & Vogan, A. A. (2022). Giant *Starship* elements mobilize accessory genes in fungal genomes. Molecular Biology and Evolution, 39(5), msac109. 10.1093/molbev/msac109

Gotoh, O., Morita, M., & Nelson, D. R. (2014). Assessment and refinement of eukaryotic gene structure prediction with gene-structure-aware multiple protein sequence alignment. BMC Bioinformatics, 15(1), 189. 10.1186/1471-2105-15-189

Grognet, P., Lalucque, H., Malagnac, F., & Silar, P. (2014). Genes that bias mendelian segregation. PLoS Genetics, 10(5), e1004387. 10.1371/journal.pgen.1004387

Gurevich, A., Saveliev, V., Vyahhi, N., & Tesler, G. (2013). QUAST: Quality assessment tool for genome assemblies. Bioinformatics, 29(8), 1072–1075. 10.1093/bioinformatics/btt086

Hall, B., Hall, M., Statisticat, L., Brown, E., Hermanson, R., Charpentier, E., Heckq, D., Laurent, S., Gronau, Q. F., & Singmann, H. (2021). Package “LaplacesDemon” (16.1.6) [Computer software]. https://github.com/LaplacesDemonR/LaplacesDemon

Hoede, C., Arnoux, S., Moisset, M., Chaumier, T., Inizan, O., Jamilloux, V., & Quesneville, H. (2014). PASTEC: An Automatic transposable element classification tool. PLoS ONE, 9(5), e91929. 10.1371/journal.pone.0091929

Hoff, K. J., Lange, S., Lomsadze, A., Borodovsky, M., & Stanke, M. (2016). BRAKER1: Unsupervised RNA-Seq-Based Genome Annotation with GeneMark-ET and AUGUSTUS. Bioinformatics, 32(5), 767–769. 10.1093/bioinformatics/btv661

Hoff, K. J., Lomsadze, A., Borodovsky, M., & Stanke, M. (2019). Whole-Genome Annotation with BRAKER. Gene Prediction, 1962, 65–95. 10.1007/978-1-4939-9173-0_5

Hymery, N., Puel, O., Tadrist, S., Canlet, C., Le Scouarnec, H., Coton, E., & Coton, M. (2017). Effect of PR toxin on THP1 and Caco-2 cells: An *in vitro* study. World Mycotoxin Journal, 10(4), 375–386. 10.3920/WMJ2017.2196

Imbernon, M., Callac, P., Gasqui, P., Kerrigan, R. W., & Velcko, A. J. (1996). *BSN*, the primary determinant of basidial spore number and reproductive mode in *Agaricus bisporus*, maps to chromosome *I*. Mycologia, 88(5), 749–761. 10.1080/00275514.1996.12026713

Iwata, H., & Gotoh, O. (2012). Benchmarking spliced alignment programs including Spaln2, an extended version of Spaln that incorporates additional species-specific features. Nucleic Acids Research, 40(20), e161–e161. 10.1093/nar/gks708

Jadhav, M. S. (2015). An update on important functionally characterized genes / QTLs of agronomic importance in crop plants. Indian Research Journal of Genetics & Biotechnology, 7(1), 44–49.

Jakubczyk, K., Kałduńska, J., Kochman, J., & Janda, K. (2020). Chemical Profile and Antioxidant Activity of the Kombucha Beverage Derived from White, Green, Black and Red Tea. Antioxidants, 9(5), 447. 10.3390/antiox9050447

Johnsson, M., Rubin, C. LJ., Höglund, A., Sahlqvist, A. LS., Jonsson, K. B., Kerje, S., Ekwall, O., Kämpe, O., Andersson, L., Jensen, P., & Wright, D. (2014). The role of pleiotropy and linkage in genes affecting a sexual ornament and bone allocation in the chicken. Molecular Ecology, 23(9), 2275–2286. 10.1111/mec.12723

Kantar, M. B., Nashoba, A. R., Anderson, J. E., Blackman, B. K., & Rieseberg, L. H. (2017). The Genetics and Genomics of Plant Domestication. BioScience, 67(11), 971–982. 10.1093/biosci/bix114

Kassambara, A., & Mundt, F. (2020). *factoextra: Extract and Visualize the Results of Multivariate Data Analyses* (R package version 1.0.7) [Computer software]. https://CRAN.R-project.org/package=factoextra

Katoh, K. (2002). MAFFT: a novel method for rapid multiple sequence alignment based on fast Fourier transform. Nucleic Acids Research, 30(14), 3059–3066. 10.1093/nar/gkf436

Kessi-Pérez, E. I., Molinet, J., & Martínez, C. (2020). Disentangling the genetic bases of *Saccharomyces cerevisiae* nitrogen consumption and adaptation to low nitrogen environments in wine fermentation. Biological Research, 53(1), 1–10. 10.1186/s40659-019-0270-3

Kieslich, P. J., Henninger, F., Wulff, D. U., Haslbeck, J. M. B., & Schulte-Mecklenbeck, M. (2019). Mouse-tracking: A practical guide to implementation and analysis. In A handbook of process tracing methods. PsyArXiv. 10.31234/osf.io/zuvqa

Kolmogorov, M., Armstrong, J., Raney, B. J., Streeter, I., Dunn, M., Yang, F., Odom, D., Flicek, P., Keane, T. M., Thybert, D., Paten, B., & Pham, S. (2018). Chromosome assembly of large and complex genomes using multiple references. Genome Research, 28(11), 1720–1732. 10.1101/gr.236273.118

Kolmogorov, M., Yuan, J., Lin, Y., & Pevzner, P. A. (2019). Assembly of long, error-prone reads using repeat graphs. Nature Biotechnology, 37(5), 540–546. 10.1038/s41587-019-00728

Kongjaimun, A., Kaga, A., Tomooka, N., Somta, P., Vaughan, D. A., & Srinives, P. (2012). The genetics of domestication of yardlong bean, *Vigna unguiculata* (L.) Walp. Ssp. Unguiculata cv.-gr. Sesquipedalis. Annals of Botany, 109(6), 1185–1200. 10.1093/aob/mcs048

Koren, S., Walenz, B. P., Berlin, K., Miller, J. R., Bergman, N. H., & Phillippy, A. M. (2017). Canu: Scalable and accurate long-read assembly via adaptive *k* -mer weighting and repeat separation. Genome Research, 27(5), 722–736. 10.1101/gr.215087.116

Krzywinski, M., Schein, J., Birol, İ., Connors, J., Gascoyne, R., Horsman, D., Jones, S. J., & Marra, M. A. (2009). Circos: An information aesthetic for comparative genomics. Genome Research, 19(9), 1639–1645. 10.1101/gr.092759.109

Kumar, J., Gupta, D. S., Gupta, S., Dubey, S., Gupta, P., & Kumar, S. (2017). Quantitative trait loci from identification to exploitation for crop improvement. Plant Cell Reports, 36(8), 1187–1213. 10.1007/s00299-017-2127-y

Kurtz, S., Phillippy, A., Delcher, A. L., Smoot, M., Shumway, M., Antonescu, C., & Salzberg, S. L. (2004). Versatile and open software for comparing large genomes. Genome Biology, 5, R12. 10.1186/gb-2004-5-2-r12

Langmead, B., & Salzberg, S. L. (2012). Fast gapped-read alignment with Bowtie 2. Nature Methods, 9(4), 357–359. 10.1038/nmeth.1923

Langmead, B., Wilks, C., Antonescu, V., & Charles, R. (2019). Scaling read aligners to hundreds of threads on general-purpose processors. Bioinformatics, 35(3), 421–432. 10.1093/bioinformatics/bty648

Laurent, B., Moinard, M., Spataro, C., Chéreau, S., Zehraoui, E., Blanc, R., Lasserre, P., Ponts, N., & Foulongne-Oriol, M. (2021). QTL mapping in *Fusarium graminearum* identified an allele of FgVe1 involved in reduced aggressiveness. Fungal Genetics and Biology, 153, 103566. 10.1016/j.fgb.2021.103566

Lê, S., Josse, J., & Husson, F. (2008). FactoMineR: An *R* Package for Multivariate Analysis. Journal of Statistical Software, 25(1). 10.18637/jss.v025.i01

Li, H. (2011). A statistical framework for SNP calling, mutation discovery, association mapping and population genetical parameter estimation from sequencing data. Bioinformatics, 27(21), 2987–2993. 10.1093/bioinformatics/btr509

Li, H. (2018). Minimap2: Pairwise alignment for nucleotide sequences. Bioinformatics, 34(18), 3094– 3100. 10.1093/bioinformatics/bty191

Liti, G., & Louis, E. J. (2012). Advances in Quantitative Trait Analysis in Yeast. PLoS Genetics, 8(8), e1002912. 10.1371/journal.pgen.1002912

Lo, Y.-C., Bruxaux, J., Rodríguez de la Vega, R. C., Snirc, A., Coton, M., Piver, M. L., Prieur, S. L., Roueyre, D., Dupont, J., Houbraken, J., Debuchy, R., Ropars, J., Giraud, T., & Branca, A. (2022). Domestication in dry-cured meat Penicillium fungi: Convergent specific phenotypes and horizontal gene transfers without strong genetic subdivision. 10.1101/2022.03.25.485132

Lomsadze, A. (2005). Gene identification in novel eukaryotic genomes by self-training algorithm. Nucleic Acids Research, 33(20), 6494–6506. 10.1093/nar/gki937

Lomsadze, A., Burns, P. D., & Borodovsky, M. (2014). Integration of mapped RNA-Seq reads into automatic training of eukaryotic gene finding algorithm. Nucleic Acids Research, 42(15), e119–e119. 10.1093/nar/gku557

Luo, R., Liu, B., Xie, Y., Li, Z., Huang, W., Yuan, J., He, G., Chen, Y., Pan, Q., Liu, Y., Tang, J., Wu, G., Zhang, H., Shi, Y., Liu, Y., Yu, C., Wang, B., Lu, Y., Han, C., … Wang, J. (2012). SOAPdenovo2: An empirically improved memory-efficient short-read *de novo* assembler. GigaScience, 1(1), 18. 10.1186/2047-217X-1-18

Maechler, M. (2016). Diptest: Hartigan’s Dip Test Statistic for Unimodality—Corrected. https://cran.r-project.org/package=diptest

Manni, M., Berkeley, M. R., Seppey, M., Simão, F. A., & Zdobnov, E. M. (2021). BUSCO Update: Novel and Streamlined Workflows along with Broader and Deeper Phylogenetic Coverage for Scoring of Eukaryotic, Prokaryotic, and Viral Genomes. Molecular Biology and Evolution, 38(10), 4647–4654. 10.1093/molbev/msab199

Marcano, Y., Montanares, M., Gil-Durán, C., González, K., Levicán, G., Vaca, I., & Chávez, R. (2023). Pr*laeA* Affects the Production of Roquefortine C, Mycophenolic Acid, and Andrastin A in Penicillium roqueforti, but It Has Little Impact on Asexual Development. Journal of Fungi, 9(10), 954. 10.3390/jof9100954

Marquina, M., Lambea, E., Carmona, M., Sánchez-Marinas, M., López-Aviles, S., Ayte, J., Hidalgo, E., & Aligue, R. (2022). A new negative feedback mechanism for MAPK pathway inactivation through Srk1 MAPKAP kinase. Scientific Reports, 12(1), 19501. 10.1038/s41598-022-23970-8

Martínez-Ainsworth, N. E., & Tenaillon, M. I. (2016). Superheroes and masterminds of plant domestication. Comptes Rendus Biologies, 339(7–8), 268–273. 10.1016/j.crvi.2016.05.005

Matas, A. J., Smith, J. M., Skeans, M. A., Thompson, B., Gustafson, S. K., Stewart, D. E., Cherikh, W. S., Wainright, J. L., Boyle, G., Snyder, J. J., Israni, A. K., & Kasiske, B. L. (2013). OPTN/SRTR 2013 Annual Data Report: Kidney. American Journal of Transplantation, 15, 1–34. 10.1111/ajt.13195

Matsuda, Y., Awakawa, T., & Abe, I. (2013). Reconstituted biosynthesis of fungal meroterpenoid andrastin A. Tetrahedron, 69(38), 8199–8204. 10.1016/j.tet.2013.07.029

McKenna, A., Hanna, M., Banks, E., Sivachenko, A., Cibulskis, K., Kernytsky, A., Garimella, K., Altshuler, D., Gabriel, S., Daly, M., & DePristo, M. A. (2010). The Genome Analysis Toolkit: A MapReduce framework for analyzing next-generation DNA sequencing data. Genome Research, 20(9), 1297–1303. 10.1101/gr.107524.110

Missiaggia, A., & Grattapaglia, D. (2006). Plant microsatellite genotyping with 4-color fluorescent detection using multiple-tailed primers. Genetics and Molecular Research, 5(1), 72–78.

Moreau, C. (1980). Le *Penicillium roqueforti*, morphologie, physiologie, intérêt en industrie fromagère, mycotoxines. Le Lait, 60(595–596), 254–271. 10.1051/lait:1980595-59615

Nguyen Ba, A. N., Lawrence, K. R., Rego-Costa, A., Gopalakrishnan, S., Temko, D., Michor, F., & Desai, M. M. (2022). Barcoded bulk QTL mapping reveals highly polygenic and epistatic architecture of complex traits in yeast. eLife, 11, e73983. 10.7554/eLife.73983

O’Gorman, C. M., Fuller, H. T., & Dyer, P. S. (2009). Discovery of a sexual cycle in the opportunistic fungal pathogen *Aspergillus fumigatus*. Nature, 457, 471–475. 10.1038/nature07528

Palmer, J., & Stajich, J. E. (2020). Funannotate v1.8.1: Eukaryotic genome annotation (1.8.1) [Computer software]. https://zenodo.org/record/4054262.

Peltier, E., Bibi-Triki, S., Dutreux, F., Caradec, C., Friedrich, A., Llorente, B., & Schacherer, J. (2021). Dissection of quantitative trait loci in the *Lachancea waltii* yeast species highlights major hotspots. G3 Genes|Genomes|Genetics, 11(9), jkab242. 10.1093/g3journal/jkab242

Petrizzelli, M., De Vienne, D., & Dillmann, C. (2019). Decoupling the Variances of Heterosis and Inbreeding Effects Is Evidenced in Yeast’s Life-History and Proteomic Traits. Genetics, 211(2), 741– 756. 10.1534/genetics.118.301635

Pfister, R., Schwarz, K. A., Janczyk, M., Dale, R., & Freeman, J. B. (2013). Good things peak in pairs: A note on the bimodality coefficient. Frontiers in Psychology, 4. 10.3389/fpsyg.2013.00700

Pinheiro, J., Bates, D., DebRoy, S., Sarkar, D., & R Core Team. (2018). {nlme}: Linear and Nonlinear Mixed Effects Models. https://cran.r-project.org/package=nlme

Punt, M., van den Brule, T., Teertstra, W. R., Dijksterhuis, J., den Besten, H. M. W., Ohm, R. A., & Wösten, H. A. B. (2020). Impact of maturation and growth temperature on cell-size distribution, heat-resistance, compatible solute composition and transcription profiles of *Penicillium roqueforti* conidia. Food Research International, 136, 109287. 10.1016/j.foodres.2020.109287

Quesneville, H., Bergman, C. M., Andrieu, O., Autard, D., Nouaud, D., Ashburner, M., & Anxolabehere, D. (2005). Combined Evidence Annotation of Transposable Elements in Genome Sequences. PLoS Computational Biology, 1(2), e22. 10.1371/journal.pcbi.0010022

Quinlan, A. R., & Hall, I. M. (2010). BEDTools: A flexible suite of utilities for comparing genomic features. Bioinformatics, 26(6), 841–842. 10.1093/bioinformatics/btq033

Rice, P., Longden, L., & Bleasby, A. (2000). EMBOSS: The European Molecular Biology Open Software Suite. Trends in Genetics, 16(6), 276–277. 10.1016/S0168-9525(00)02024-2

Rojas-Aedo, J. F., Gil-Durán, C., Goity, A., Vaca, I., Levicán, G., Larrondo, L. F., & Chávez, R. (2018). The developmental regulator Pcz1 affects the production of secondary metabolites in the filamentous fungus *Penicillium roqueforti*. Microbiological Research, 212–213, 67–74. 10.1016/j.micres.2018.05.005

Ropars, J., López-Villavicencio, M., Dupont, J., Snirc, A., Gillot, G., Coton, M., Jany, J.-L., Coton, E., & Giraud, T. (2014). Induction of sexual reproduction and genetic diversity in the cheese fungus *Penicillium roqueforti*. Evolutionary Applications, 7(4), 433–441. 10.1111/eva.12140

Ropars, J., Rodríguez de la Vega, R. C., López-Villavicencio, M., Gouzy, J., Sallet, E., Dumas, É., Lacoste, S., Debuchy, R., Dupont, J., Branca, A., & Giraud, T. (2015). Adaptive Horizontal Gene Transfers Between Multiple Cheese-Associated Fungi. Current Biology, 25(19), 2562–2569. 10.1016/j.cub.2015.08.025

Ropars, J., Toro, K. S., Noel, J., Pelin, A., Charron, P., Farinelli, L., Marton, T., Krüger, M., Fuchs, J., Brachmann, A., & Corradi, N. (2016). Evidence for the sexual origin of heterokaryosis in arbuscular mycorrhizal fungi. Nature Microbiology, 1(6), 1–9. 10.1038/nmicrobiol.2016.33

Ropars, J., Caron, T., Lo, Y.-C., Bennetot, B., & Giraud, T. (2020). The domestication of Penicillium cheese fungi. Comptes Rendus - Biologies, 1(0). 10.5802/crbiol.15

Ropars, J., & Giraud, T. (2022). Convergence in domesticated fungi used for cheese and dry-cured meat maturation: Beneficial traits, genomic mechanisms, and degeneration. Current Opinion in Microbiology, 70, 102236. 10.1016/j.mib.2022.102236

Schneider, C. A., Rasband, W. S., & Eliceiri, K. W. (2012). NIH Image to ImageJ: 25 years of image analysis. Nature Methods, 9(7), 671–675. 10.1038/nmeth.2089

Schuelke, M. (2000). An economic method for the fluorescent labeling of PCR fragments. A poor man’s approach to genotyping for research and high-throughput diagnostics. Nature Biotechnology, 18, 233–234. 10.1038/72708

Scott, P. M. (1981). Toxins of *Penicillium* Species Used in Cheese Manufacture. Journal of Food Protection, 44(9), 702–710. 10.4315/0362-028X-44.9.702

Seekles, S. J., Teunisse, P. P. P., Punt, M., Van Den Brule, T., Dijksterhuis, J., Houbraken, J., Wösten, H. A. B., & Ram, A. F. J. (2021). Preservation stress resistance of melanin deficient conidia from *Paecilomyces variotii* and *Penicillium roqueforti* mutants generated via CRISPR/Cas9 genome editing. Fungal Biology and Biotechnology, 8(1), 4. 10.1186/s40694-021-00111-w

Silva, M. V. B., dos Santos, D. J. A., Boison, S. A., Utsunomiya, A. T. H., Carmo, A. S., Sonstegard, T. S., Cole, J. B., & Van Tassell, C. P. (2014). The development of genomics applied to dairy breeding. Livestock Science, 166(1), 66–75. 10.1016/j.livsci.2014.05.017

Smit, A., & Hubley, R. (2008). *RepeatModeler Open-1.0* (1.0) [Computer software]. http://www.repeatmasker.org

Smit, A., Hubley, R., & Green, P. (2013). *RepeatMasker Open-4.0.* (4.0) [Computer software]. http://www.repeatmasker.org

Solares, E. A., Chakraborty, M., Miller, D. E., Kalsow, S., Hall, K., Perera, A. G., Emerson, J. J., & Hawley, R. S. (2018). Rapid Low-Cost Assembly of the *Drosophila melanogaster* Reference Genome Using Low-Coverage, Long-Read Sequencing. G3 Genes|Genomes|Genetics, 8(10), 3143–3154. 10.1534/g3.118.200162

Somta, P., Chen, J., Yimram, T., Yundaeng, C., Yuan, X., Tomooka, N., & Chen, X. (2020). QTL Mapping for Agronomic and Adaptive Traits Confirmed Pleiotropic Effect of *mog* Gene in Black Gram [*Vigna mungo* (L.) Hepper]. Frontiers in Genetics, 11, 635. 10.3389/fgene.2020.00635

Stanke, M., Schöffmann, O., Morgenstern, B., & Waack, S. (2006). Gene prediction in eukaryotes with a generalized hidden Markov model that uses hints from external sources. BMC Bioinformatics, 7(1), 62. 10.1186/1471-2105-7-62

Stanke, M., Diekhans, M., Baertsch, R., & Haussler, D. (2008). Using native and syntenically mapped cDNA alignments to improve de novo gene finding. Bioinformatics, 24(5), 637–644. 10.1093/bioinformatics/btn013

Steinmetz, L. M., Sinha, H., Richards, D. R., Spiegelman, J. I., Oefner, P. J., McCusker, J. H., & Davis, R. W. (2002). Dissecting the architecture of a quantitative trait locus in yeast. Nature, 416(6878), 326–330. 10.1038/416326a

Sweeney, M., & McCouch, S. (2007). The Complex History of the Domestication of Rice. Annals of Botany, 100(5), 951–957. 10.1093/aob/mcm128

Swinnen, S., Thevelein, J. M., & Nevoigt, E. (2012). Genetic mapping of quantitative phenotypic traits in *Saccharomyces cerevisiae*. FEMS Yeast Research, 12(2), 215–227. 10.1111/j.1567-1364.2011.00777.x

Tang, H., Zhang, X., Miao, C., Zhang, J., Ming, R., Schnable, J. C., Schnable, P. S., Lyons, E., & Lu, J. (2015). ALLMAPS: Robust scaffold ordering based on multiple maps. Genome Biology, 16(1), 3. 10.1186/s13059-014-0573-1

Taylor, J., & Butler, D. (2017). *R* Package ASMap: Efficient Genetic Linkage Map Construction and Diagnosis. Journal of Statistical Software, 79(6). 10.18637/jss.v079.i06

Telias, A., Lin-Wang, K., Stevenson, D. E., Cooney, J. M., Hellens, R. P., Allan, A. C., Hoover, E. E., & Bradeen, J. M. (2011). Apple skin patterning is associated with differential expression of *MYB10*. BMC Plant Biology, 11(1), 93. 10.1186/1471-2229-11-93

Todd, R. T., Forche, A., & Selmecki, A. (2017). Ploidy Variation in Fungi—Polyploidy, Aneuploidy, and Genome Evolution. Microbiology Spectrum, 5(4), 1–31. 10.1128/microbiolspec.FUNK-0051-2016.

Torrent, C., Gil-Durán, C., Rojas-Aedo, J. F., Medina, E., Vaca, I., Castro, P., García-Rico, R. O., Cotoras, M., Mendoza, L., Levicán, G., & Chávez, R. (2017). Role of *sfk1* Gene in the Filamentous Fungus *Penicillium roqueforti*. Frontiers in Microbiology, 8, 2424. 10.3389/fmicb.2017.02424

Turner, B. C., & Perkins, D. D. (1979). SPORE KILLER, A CHROMOSOMAL FACTOR IN NEUROSPORA THAT KILLS MEIOTIC PRODUCTS NOT CONTAINING IT. Genetics, 93(3), 587–606. 10.1093/genetics/93.3.587

Untergasser, A., Cutcutache, I., Koressaar, T., Ye, J., Faircloth, B. C., Remm, M., & Rozen, S. G. (2012). Primer3-new capabilities and interfaces. Nucleic Acids Research, 40(15), e115. 10.1093/nar/gks596

Van Wyk, S., Harrison, C. H., Wingfield, B. D., De Vos, L., Van Der Merwe, N. A., & Steenkamp, E. T. (2019). The RIPper, a web-based tool for genome-wide quantification of Repeat-Induced Point (RIP) mutations. PeerJ, 7, e7447. 10.7717/peerj.7447

Vernozy-Rozand, C., Mazuy-Cruchaudet, C., Bavai, C., Montet, M. P., Bonin, V., Dernburg, A., & Richard, Y. (2005). Growth and survival of *Escherichia coli* O157:H7 during the manufacture and ripening of raw goat milk lactic cheeses. International Journal of Food Microbiology, 105(1), 83–88. 10.1016/j.ijfoodmicro.2005.05.005

Vion, C., Peltier, E., Bernard, M., Muro, M., & Marullo, P. (2021). Marker Assisted Selection of Malic-Consuming Saccharomyces cerevisiae Strains for Winemaking. Efficiency and Limits of a QTL’s Driven Breeding Program. Journal of Fungi, 7(4), 304. 10.3390/jof7040304

Walker, B. J., Abeel, T., Shea, T., Priest, M., Abouelliel, A., Sakthikumar, S., Cuomo, C. A., Zeng, Q., Wortman, J., Young, S. K., & Earl, A. M. (2014). Pilon: An Integrated Tool for Comprehensive Microbial Variant Detection and Genome Assembly Improvement. PLoS ONE, 9(11), e112963. 10.1371/journal.pone.0112963

Wang, J., Liao, X., Li, Y., Zhou, R., Yang, X., Gao, L., & Jia, J. (2010). Fine mapping a domestication-related QTL for spike-related traits in a synthetic wheat. Genome, 53(10), 798–804. 10.1139/G10-066

Wei, T., & Simko, V. (2021). *R package “corrplot”: Visualization of a Correlation Matrix* (0.92) [Computer software]. https://github.com/taiyun/corrplot

Wick, R. R., Judd, L. M., Gorrie, C. L., & Holt, K. E. (2017). Completing bacterial genome assemblies with multiplex MinION sequencing. Microbial Genomics, 3(10). 10.1099/mgen.0.000132

Wicker, T., Sabot, F., Hua-Van, A., Bennetzen, J. L., Capy, P., Chalhoub, B., Flavell, A., Leroy, P., Morgante, M., Panaud, O., Paux, E., SanMiguel, P., & Schulman, A. H. (2007). A unified classification system for eukaryotic transposable elements. Nature Reviews Genetics, 8(12), 973–982. 10.1038/nrg2165

Wickham, H. (2007). Reshaping Data with the {reshape} Package. Journal of Statistical Software, 21(12), 1–20.

Wickham, H. (2016). *ggplot2: Elegant Graphics for Data Analysis* [Computer software]. https://ggplot2.tidyverse.org

Wilkening, S., Lin, G., Fritsch, E. S., Tekkedil, M. M., Anders, S., Kuehn, R., Nguyen, M., Aiyar, R. S., Proctor, M., Sakhanenko, N. A., Galas, D. J., Gagneur, J., Deutschbauer, A., & Steinmetz, L. M. (2014). An Evaluation of High-Throughput Approaches to Qtl Mapping in *Saccharomyces Cerevisiae*. Genetics, 196(3), 853–865. 10.1534/genetics.113.160291

Wirén, A., & Jensen, P. (2011). A Growth QTL on Chicken Chromosome 1 Affects Emotionality and Sociality. Behavior Genetics, 41(2), 303–311. 10.1007/s10519-010-9377-6

Wright, D. (2015). The Genetic Architecture of Domestication in Animals. Bioinformatics and Biology Insights, 9S4, BBI.S28902. 10.4137/BBI.S28902

Wright, D., Rubin, C.-J., Martinez Barrio, A., Schütz, K., Kerje, S., Brändström, H., Kindmark, A., Jensen, P., & Andersson, L. (2010). The genetic architecture of domestication in the chicken: Effects of pleiotropy and linkage. Molecular Ecology, 19(23), 5140–5156. 10.1111/j.1365-294X.2010.04882.x

Yao, J., Xu, J., Cornille, A., Tomes, S., Karunairetnam, S., Luo, Z., Bassett, H., Whitworth, C., ReesLGeorge, J., Ranatunga, C., Snirc, A., Crowhurst, R., De Silva, N., Warren, B., Deng, C., Kumar, S., Chagné, D., Bus, V. G. M., Volz, R. K., … Gleave, A. P. (2015). A *micro RNA* allele that emerged prior to apple domestication may underlie fruit size evolution. The Plant Journal, 84(2), 417–427. 10.1111/tpj.13021

Zimmer, A., Durand, C., Loira, N., Durrens, P., Sherman, D. J., & Marullo, P. (2014). QTL Dissection of Lag Phase in Wine Fermentation Reveals a New Translocation Responsible for *Saccharomyces cerevisiae* Adaptation to Sulfite. PLoS ONE, 9(1), e86298. 10.1371/journal.pone.0086298

