## Supplementary files containing Supplementary figures 1 to 6 for "Generation of diversity in the blue cheese mold *Penicillium roqueforti* and identification of pleiotropic QTL for key cheese-making phenotypes"

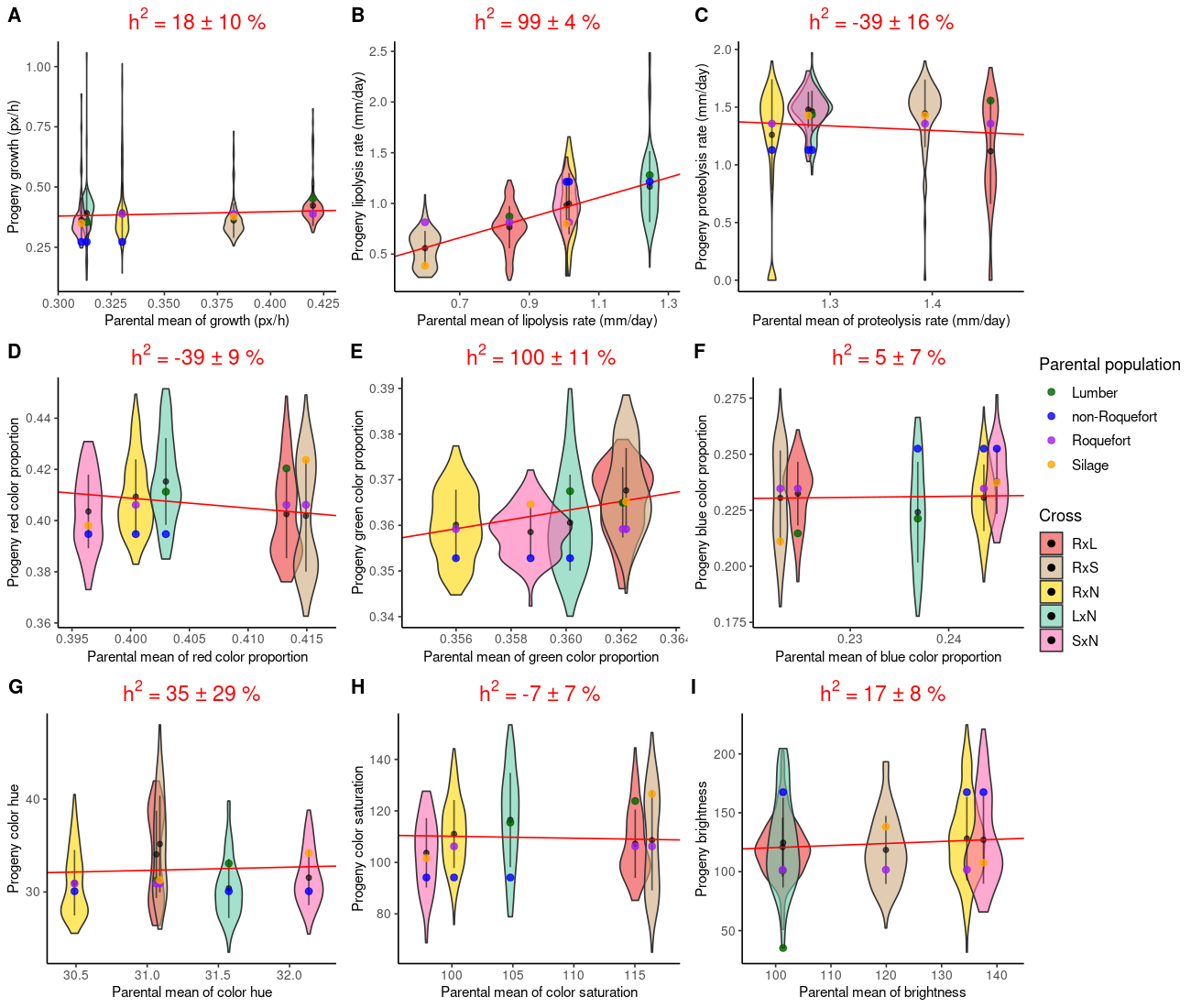

Supplementary Figure 1: Distributions of phenotypes measured in the five progenies as a function of the parental means: (A) diametral growth in pixel per hour, (B) lipolysis rate in mm per day, (C) proteolysis rate in mm per day, (D) relative red, (E) green, (F) blue, (G) hue, (H) saturation and (I) brightness; progenies are represented along the x axis, RxL in red, RxS in beige, RxN in yellow, LxN in turquoise and SxN in pink. The black point and line represent the mean and standard deviation, respectively, in progenies. The coloured points represent the parental values (lumber in green, non-Roquefort in blue, Roquefort in purple, silage in orange). The red line represents the linear regression from which the slope is used to estimate the heritability value (h^2^) given at the top.

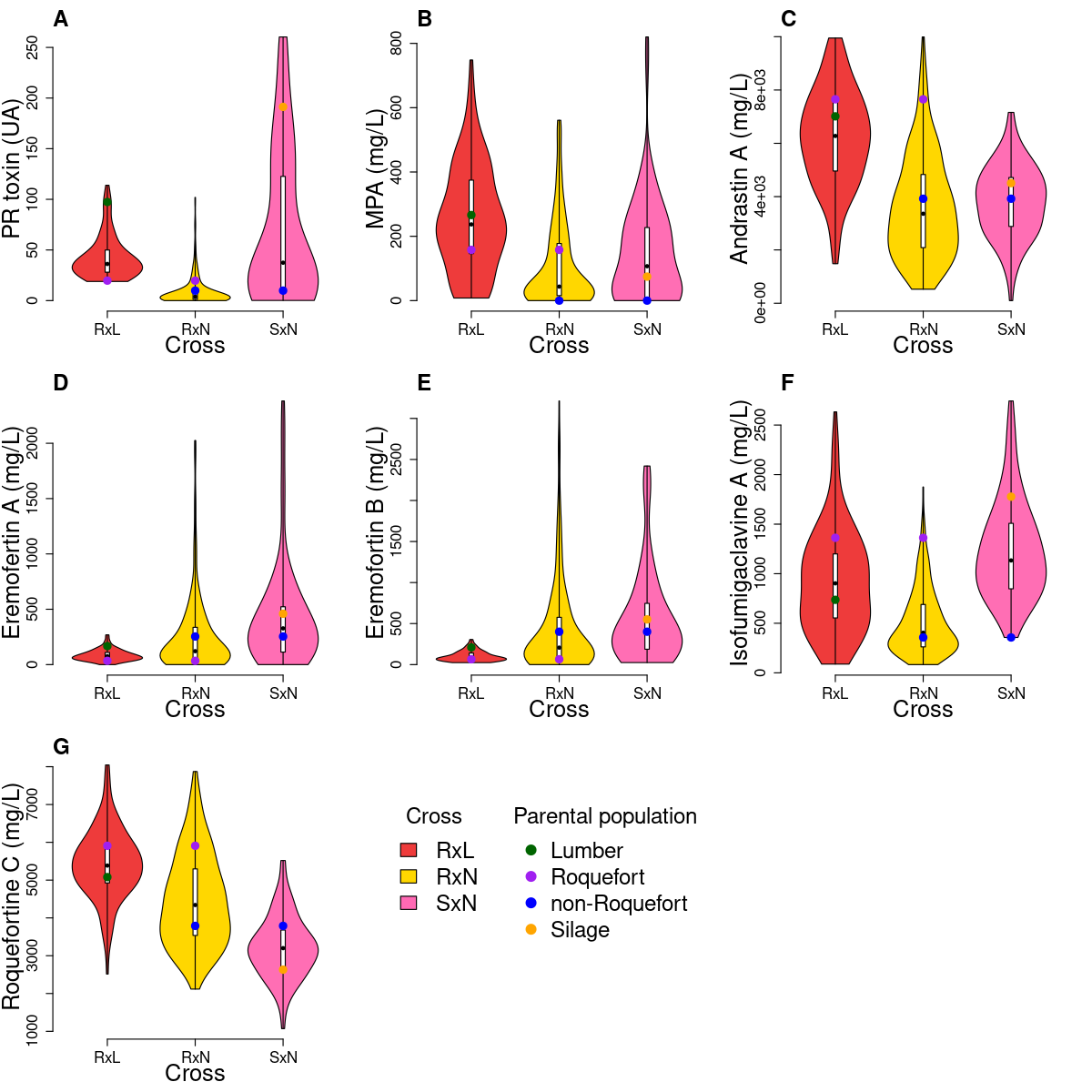

Supplementary Figure 2: Distributions of extrolite production in three progenies: (A) PR toxin, (B) mycophenolic acid, (C) andrastin A, (D) eremofortin A, (E) eremofortin B, (F) (iso)-fumigaclavine A and (G) roquefortine C; progenies are represented along the x axis: RxL in red, RxN in yellow and SxN in pink. The coloured points represent the parental values (lumber in green, non-Roquefort in blue, Roquefort in purple, silage in orange). All units are in mL.L^-1^ except the PR toxin, which is expressed in arbitrary units.

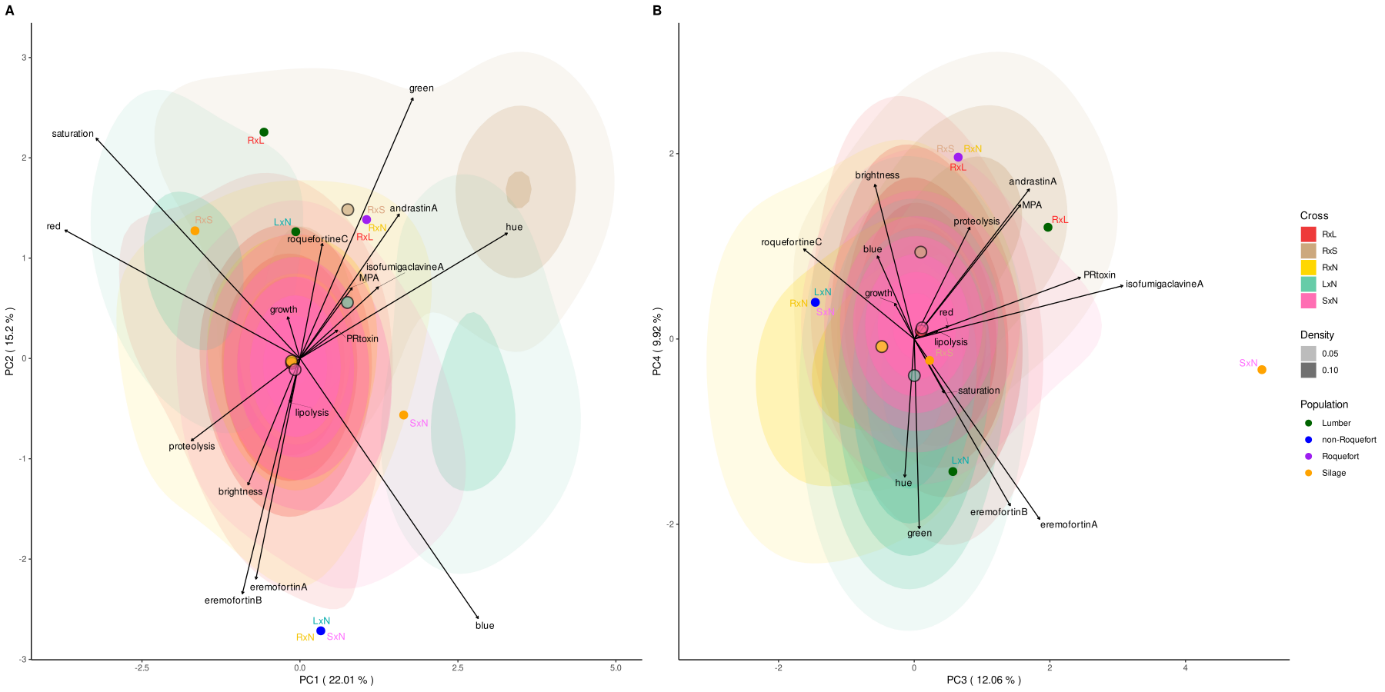

Supplementary Figure 3: Biplot of a principal component analysis with the two first components (A) and the third and fourth (B). For every cross the progeny scores are displayed in a different color (LxN in red, RxL in brown, RxN in yellow, RxS in turquoise, SxN in pink) and with two density levels (from 0.05 to 0.10 in light color and above 0.10 in dark color). Large dots represent the barycentre of each progeny’s scores using the same color code (the red one is under the yellow one). The 4-fold loadings of the phenotypes are represented by the black arrows. Small dots show the scores of the parental strains from different populations (lumber in green, non-Roquefort in blue, Roquefort in purple and silage in orange) in the labels show the cross they are involved in. The percentage in the axis labels shows the total variance explained by the component.

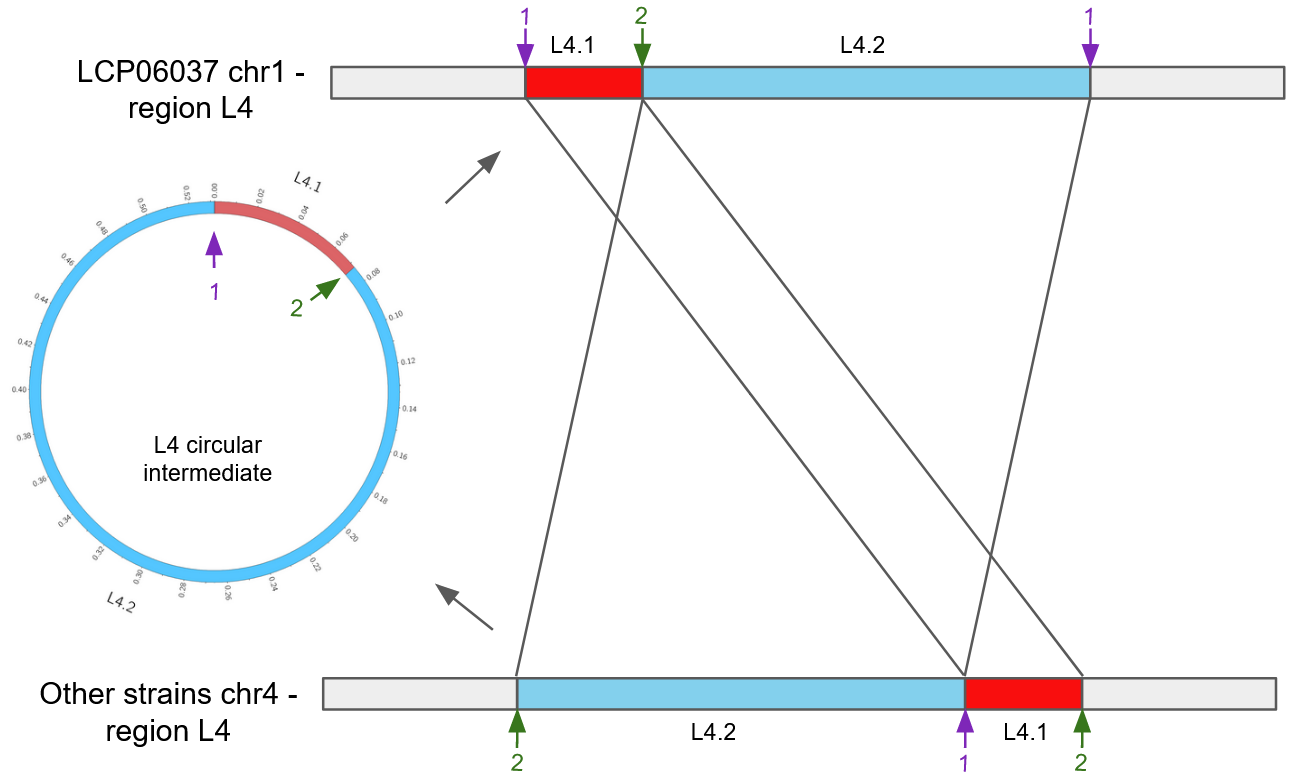

Supplementary Figure 4: Focus on the L4 translocated region in the *Penicillium roqueforti* LCP06037 strain, the same pattern being observed for the L1 region in the LCP06039 strain. The L4 region is subdivided into two subregions, L4.1 in red and L4.2 in blue, with different relative positions, suggesting a circular intermediate for the translocation. The cut sites for the proposed circular intermediate are indicated by purple and green arrows, respectively, and numbered 1 and 2. The figure shows the synteny of the L4 translocated region present in the chromosome 1 in the LCP06037 lumber parental strain and in the chromosome 4 in other parents and the proposed circular intermediate.

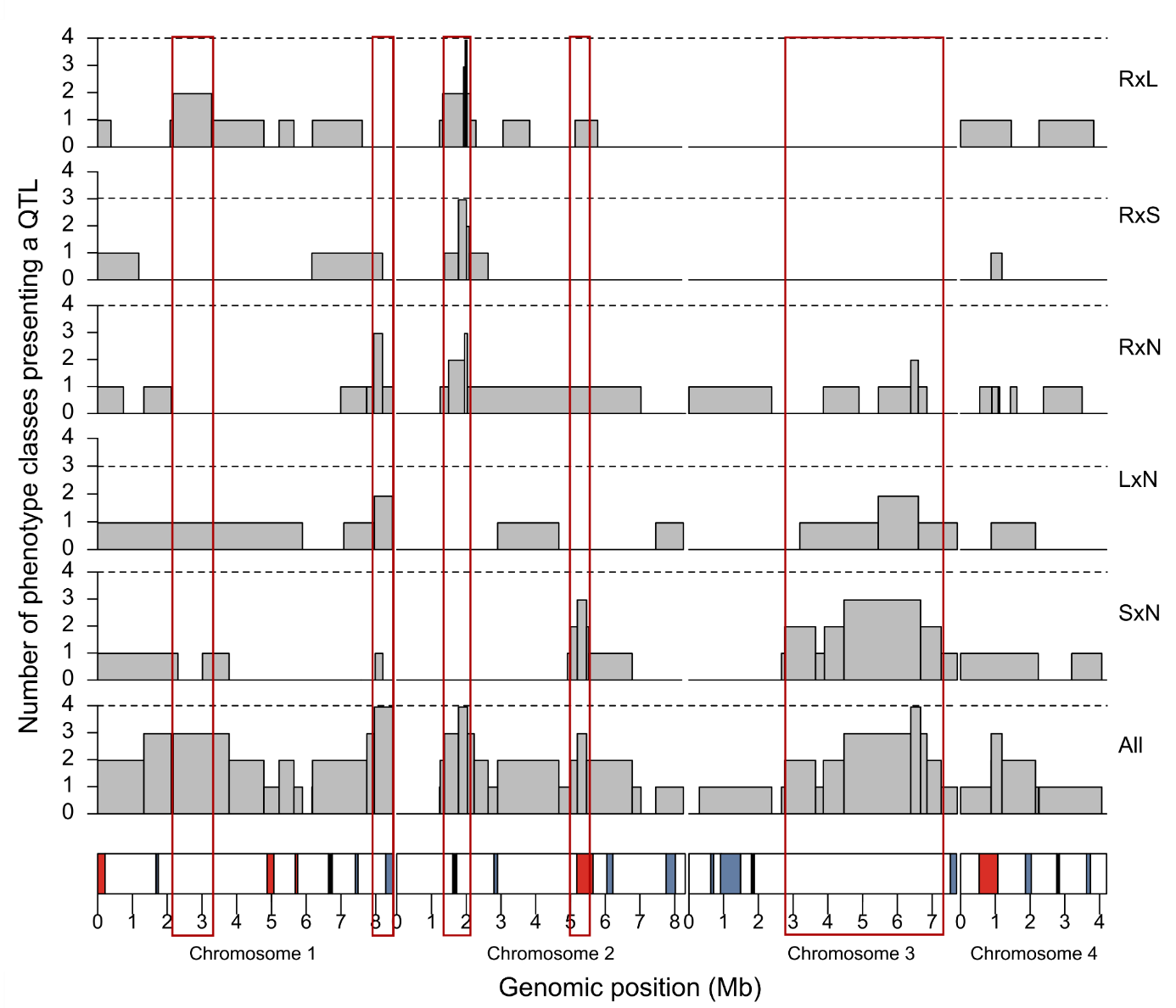

Supplementary Figure 5: Number of phenotype classes (*i.e.* lipolysis, proteolysis, color and extrolite production) presenting a QTL along the genome positions. The crosses are represented one per line, with the cross ID indicated on the right, the first parent carrying the MAT1-1 mating type; the bottom line represents a pool of all crosses. The x-axis represents genomic physical positions (Mb). At the bottom, the four chromosomes of the reference genome (LCP06133) are represented by rectangles depicting genomic features: large horizontally transferred regions specific to the reference genome, in blue, and translocated regions in the reference genome, in red. The large vertical empty red rectangles indicate pleiotropic QTL regions, with effects on multiple traits in at least one cross.

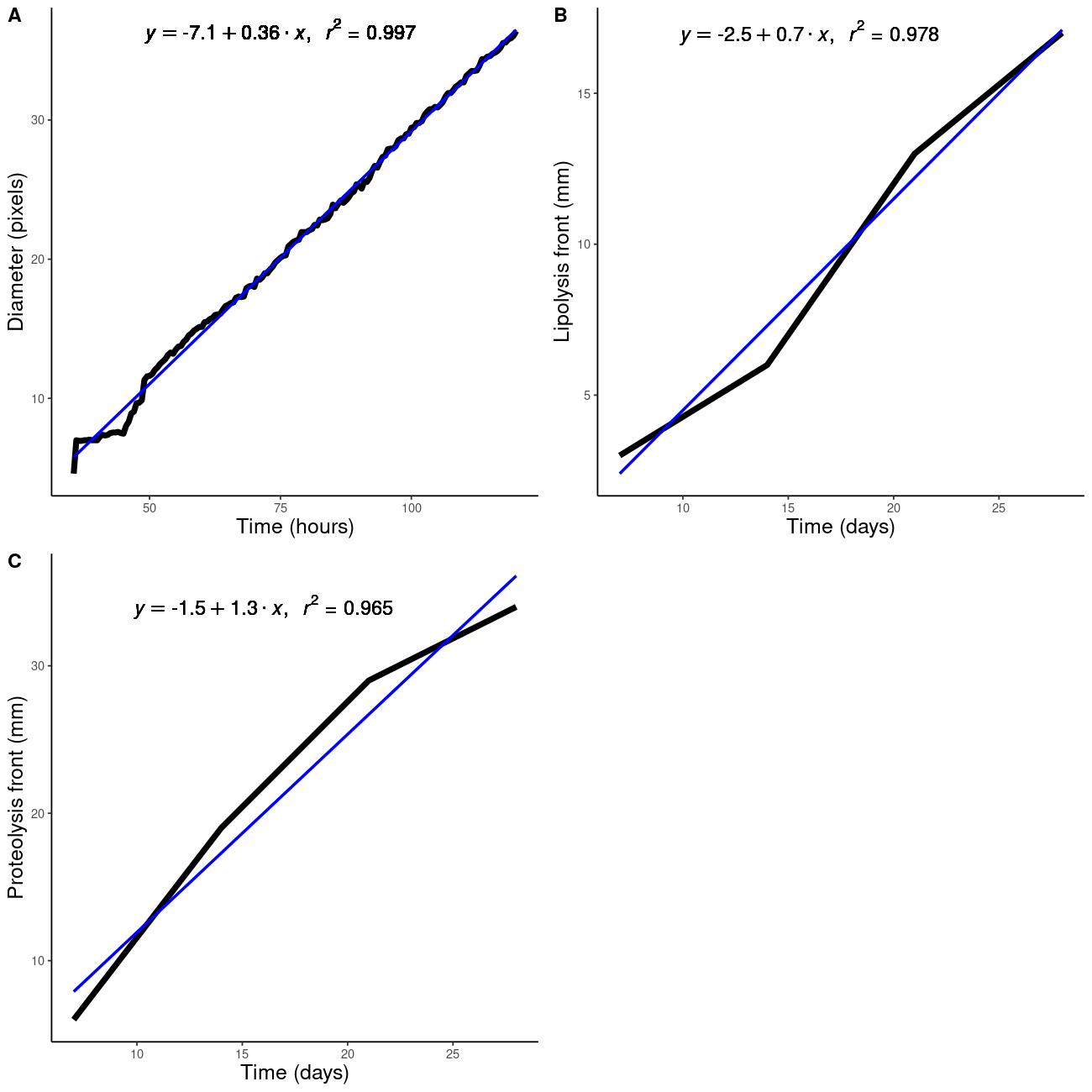

Supplementary Figure 6: Example of phenotype measures and parameter estimation, for an offspring having a median r^2^ value for the least squares regression fit: (A) growth dynamic, *i.e.* colony diameter (in pixel numbers on pictures) as a function of time (hours), (B) lipolysis dynamics and (C) proteolysis dynamics, B and C representing the distance of the lysis front (*i.e* characterized by medium fading) in mm per day. The linear regression lines are shown in blue, and their equations and coefficients of determination are given at the top.
